# Interactions between hippocampus and visceral organs in sleep and wakefulness

**DOI:** 10.64898/2026.04.28.721215

**Authors:** Ekaterina Levichkina, Ivan N. Pigarev, Trichur R. Vidyasagar

**Author notes:** Deceased (Ivan N. Pigarev). Correspondence should be addressed to Trichur Vidyasagar.

## Abstract

Exteroception is attenuated during sleep, while interoceptive signals may remain active. If so, they can trigger responses in their target areas, including the hippocampus, which is known to receive information from the internal organs and is susceptible to oversynchronization while triggered. We investigated whether the hippocampus is synchronized to various visceral events in sleep and wakefulness.

Activity of hippocampal neurons and local field potentials (LFPs) was co-registered with respiration, heart rate and myoelectric signals of the stomach and duodenum in two adult female cats over multiple sleep–wake cycles. Visceral event-triggered and neuronal spike-triggered (bootstrapping-based) analyses were performed in wakefulness and SWS.

Synchronization between visceral and hippocampal activities occurred in both wakefulness and SWS. However, hippocampal cells and LFPs showed preferences for one state only. Consistent with prior studies, we found the strongest link between high-amplitude respiratory events and hippocampal activity, with significantly higher occurrence during SWS. Both stomach and duodenal signals were also represented in hippocampal activity. Motility-associated duodenal myoelectric signals correlated with hippocampal activity more during wakefulness where gastrointestinal motility is more active, while synchronization between regular duodenal waves and the hippocampus was more frequent during SWS.

We conclude that the interoceptive signals reach the hippocampus in both sleep and wakefulness and suggest that they have the potential to oversynchronize any ongoing synchronized slow-wave activities in the hippocampal network during slow wave sleep (SWS).

**Significance Statement:** What types of sensory input shapes hippocampal activity and regulates its rhythmical structure is important for understanding both normal and paroxysmal hippocampal dynamics. We find that synchronization between interoceptive signals and hippocampal neural activity exists during wakefulness and is preserved during SWS. Moreover, in some cases, this association increases in SWS for respiratory and periodic duodenal activities. Our data revealed specificity of hippocampal neurons to both type of visceral activity and state of vigilance, suggesting that hippocampal network is shaped by visceral activity in a state-dependent manner. Our results highlight the factors underlying the comorbidities of epilepsy with gastrointestinal, cardiac or respiratory disorders.

## Introduction

Hippocampal activity is known to be strongly influenced by the context of concurrently occurring events (Vidyasagar et al., 1991; Kubie and Muller, 1991; Salzmann et al., 1993; Kennedy and Shapiro, 2009; Preston et al., 2013; Azevedo et al., 2019). This is related to various aspects of the external environment, such as odours or visual cues or to internal signals reflecting an animal’s emotional state or current needs. The hippocampus is ideally placed for assessing emotional states and modulating internal goals due to its connections to brain structures involved in visceral or emotional control. These include hypothalamic zones associated with feeding, reproduction and defence mechanisms, the insular cortex crucial for viscerosensory and visceromotor functions, the infralimbic and prelimbic cortices and the amygdala (Kishi et al., 2000; Petrovich et al., 2001; Castle et al., 2005; Herman et al., 2005; Dong and Swanson, 2006; Fanselow and Dong, 2010).

Hippocampal neurons respond to interoceptive signals related to breathing and heart rate (Frysinger and Harper, 1988). Synchronization of hippocampal activity with breathing has been reported in various species (Duffin and Hockman, 1972; Frysinger et al., 1989; Radna and MacLean, 1981; Poe et al., 1996; Herrero et al., 2018; Folschweiller and Sauer, 2021; Nokia and Penttonen, 2022). However, much less is known about the hippocampal role in analysing interoceptive information from other modalities. Given the hippocampus’s connections to diverse brain regions involved in visceral processing, such relationships warrant investigation.

The context-dependency of hippocampal activity is typically discussed for awake behaviour and exteroceptive signals, well documented in macaques across routine situations (Vidyasagar et al., 1991; Salzman et al., 1993). However, signals related to visceral activity are also available to the brain, especially during sleep, since certain visceral processes are specifically associated with sleep (e.g., Moore, 1991; Besedovsky et al., 2012; Ali et al., 2013). Emerging evidence suggests that the brain may be more receptive to internal signals during sleep when behavioural distractions and inhibitory sympathetic signals are attenuated (Pigarev, 1994; Pigarev et al., 2013; Rembado et al., 2021; Levichkina et al., 2021, 2022).

Across species, from *Drosophila* to humans, satiety has been linked to sleep (Borbély, 1977; Orr et al., 1997; Bazar, Yun, and Lee, 2004; Murphy et al., 2016). Postprandial sleepiness has been shown in rodent studies to result from the production of cholecystokinin and other gastrointestinal peptides in the digestive tract (Shemyakin and Kapás, 2001; Kim and Lee, 2009). Recent studies demonstrated marked synchronization between stomach and brain activities during resting state, involving multiple brain areas including the limbic system (Cao et al., 2022; Rebollo et al., 2018). This suggests that certain digestive processes predominantly occur during rest, with interoceptive signals reaching their brain targets mainly in this state (Levichkina et al., 2021).

The hippocampus is known to be highly susceptible to seizures, including those provoked by external and internal events (Okudan et al., 2018; Girges et al., 2020; Hanif and Musick, 2021; Strzelecka et al., 2022). Considering that multiple types of epileptiform activity occur more frequently during SWS or drowsiness compared to wakefulness (Shouse et al., 1996; Herman et al., 2001; Dinner, 2002; Combi et al., 2004; Pavlova et al., 2004; Hofstra and de Weerd, 2009; Chokroverty and Nobili, 2017; Garg et al., 2022) and are enhanced during SWS in mesial temporal epilepsy (Nazer and Dickson, 2009), it seems essential to study the causes of synchronization of hippocampal activity across the sleep-wake cycle. If uninhibited or amplified interoceptive signals are transmitted to the hippocampus during sleep, they may act as triggers for oversynchronization (Levichkina et al., 2025). Although the hippocampus receives visceral signals in wakefulness, they may have an enhanced potential to oversynchronize hippocampal networks during SWS due to the potentially higher degree of brain synchronization within the hippocampus during SWS.

We investigated whether activity of hippocampal cells and local field potentials synchronized with various visceral events and whether they do so differently between sleep and wakefulness.

## Materials and Methods

### EXPERIMENTAL DESIGN

The Russian foundations distributing scientific funding conduct an ethics evaluation of every research proposal prior to making a decision regarding financial support for a particular study. This evaluation is performed by the Council of Reviewers. Upon receiving funding support from the Russian Foundation for Basic Research (RFBR), an ethical assessment of the presented research was conducted also by the Institutional Scientific Councils of IITP Russian Academy of Sciences, guided by the Ethical Principles for the Maintenance and Use of Animals in Neuroscience Research (Zimmermann, 1987).

The experiments were conducted on two adult female cats, with neuronal activity in the hippocampus recorded via painless head fixation using a halo implant (for details of the method, see Pigarev et al., 2009; Levichkina et al., 2021). Recordings were performed across several natural sleep-wake cycles, with each session lasting up to four hours per day.

Several parameters reflecting visceral activity were co-registered with hippocampal activity. Besides electrocardiogram (ECG), breathing rate (BR), myoelectric intestinal activity and myoelectric gastric activity, electroencephalogram (EEG) recordings and eye movement data were collected to assist in identifying the state of vigilance. Figure 1 schematically illustrates the recorded parameters. Panel A depicts the approach to hippocampal activity registration, panel B shows the co-registered signals, and panel C provides a 30-second example of the recorded data.

**Figure 1.**
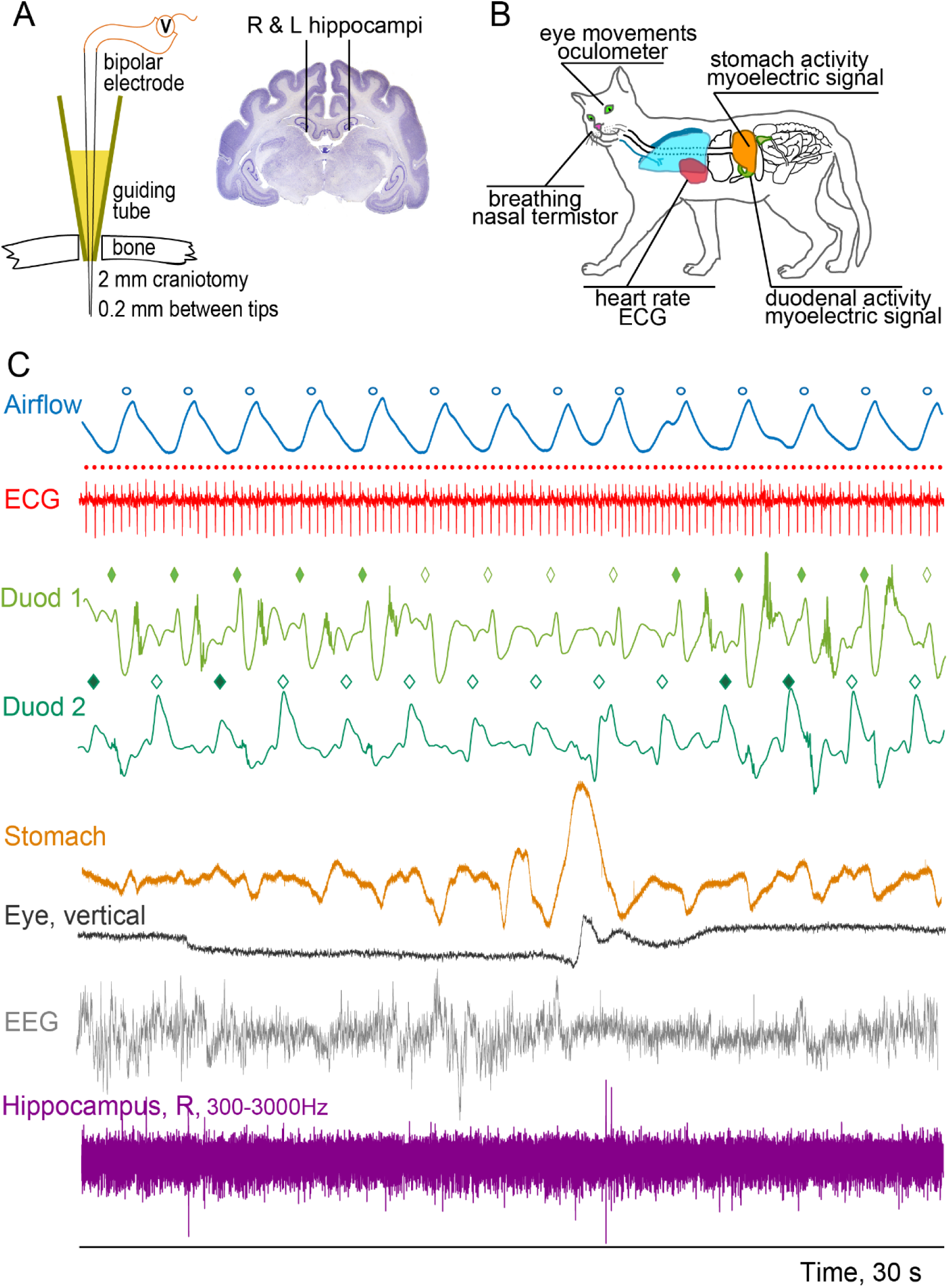
Data acquisition and types of recorded signals. A. Approach for recording brain activity: bipolar electrode consisting of 2 etched and varnished tungsten wires, introduced into the dorsal hippocampus with the help of a conical guide tube. B. Schematic representation of the sources of recorded signals. Modified from: Amsel, Sheri. “Cat Anatomy (Thoracic and Abdominal Organs)” Exploring Nature Educational Resource ©2005-2024. March 25, 2024 <http://www.exploringnature.org/db/view/Cat-Anatomy-Thoracic-and-Abdominal-Organs > C. 30-seconds long example of all co-recorded activities and markers of periodic events. Markers are shown above their corresponding signals: air flow markers (open blue dots), heart beats (filled red dots), and 2 duodenal myoelectric signals (Duod) recorded from 2 different parts of the duodenal wall (diamonds). Open diamonds signify SW periods, and filled diamonds show SB periods. One can see that different parts of duodenum are only partially synchronized. The episode of data shows an interval of slow wave sleep at the beginning (note high EEG amplitude in grey) with nearly absent stomach activity, followed by stomach contraction accompanied by several events: decreased SWS in the EEG, brief eye movement (black), heart rate modulation (red) and firing of some hippocampal cells (violet).

The animals were habituated to the laboratory environment through a process commonly referred to as “shaping,” which lasted 2 to 3 weeks and was based on positive reinforcement. During this time, the animals were fed in the lab space, including on top of the recording table, and were allowed to freely explore the laboratory space for at least two hours per day. Once shaping was successfully completed, the cats underwent MRI scanning, using a 1.5 Tesla magnet. Such pre-surgical MRI is essential for identifying individual stereotaxic coordinates for electrode placement in chronic experiments, where histological verification of each electrode track is not feasible. The technique for MRI-based verification of stereotaxic coordinates in cats was described in detail by Levichkina et al. (2021).

The surgery was conducted in two stages. The first surgical procedure involved attaching a “halo” head frame to the skull, placing EEG electrodes and creating small craniotomies over the two hippocampi and a guiding conical tube inserted above the dura for later electrode placements. In the second procedure, recording electrodes for registering myoelectric activity from the duodenal and gastric walls were implanted through the guide tubes. See Pigarev et al. (2009) for details of this technique.

All surgeries were performed under general anaesthesia. Xylazine (0.5 mg/kg, IM) was administered as premedication and for analgesia. Anaesthesia was induced and maintained with Zoletil, beginning with an initial injection of 6 mg/kg. Additional 5 mg doses were given as needed to sustain surgical levels of anaesthesia, using changes in respiration and heart rates to administer the top-ups to prevent paw withdrawal reflex. For the two cats used in this study, the frequency of the additional doses needed was approximately every 20 minutes. Each surgery lasted less than two hours.

### ATTACHMENT OF “HALO”

This implant was used for painless head fixation during recordings to reduce motion artefacts and ensure stable data quality (see Pigarev et al., 2009, for a detailed description of the techniques, and Levichkina et al., 2021, for their application in cats). After anesthetizing the animal and placing its head in the stereotaxic frame, the soft tissues were removed from the dorsal surface of its skull. Eight small orthopaedic screws were inserted into the skull without penetrating the dura, approximately 15 mm from the sagittal suture on both the left and right sides. These screws were connected by stainless steel wire to serve as the base of the halo frame. The frame was reinforced with dental acrylic cement, which was also used to create a thin covering layer on the skull surface.

A pair of EEG screw electrodes made of Elgiloy were screwed in through 2 mm craniotomies above the occipital and frontal lobes and covered with acrylic cement. Stereotaxic coordinates were transferred onto the transparent covering layer, and one or two 2 mm craniotomies were made at the coordinates above the hippocampus. These craniotomies were covered with sterile conical tubes filled with sterile bone wax to serve as guiding tubes for electrode penetrations. This technique permitted multiple electrode penetrations through the same craniotomy and allowed for easy repositioning of the craniotomy to change the recording site if necessary, as 2 mm craniotomies healed quickly after the guiding tube’s removal.

The halo enabled painless head fixation while allowing sufficient body movement, so the cat could assume a comfortable position and change it at will, apart from the head position. Training for this started two-weeks after the surgery. The training duration (maximum of 4 hours) was guided by the cat’s acceptance and usually took 2–3 weeks, at which point the animal began to sleep with its head fixed, as required for the planned experiments. It has been noticed in experiments on cats in Moscow by author Pigarev and on macaques in Melbourne by all three authors that during periods when the animals were not engaged in a behavioural task, they often tend to fall asleep. This is likely due to the relaxation of the neck muscles, since the muscles are relieved of the weight of the head by the head fixation.

Recordings were made from the right hippocampus in cat 1 at the following coordinates: AP +2 mm, LAT 9 mm. In cat 2, recordings were made from both left and right hippocampi at AP +5 mm, LAT 5 mm, and AP +6 mm, LAT 3.5 mm, respectively. In both animals, the recording depth ranged from 11 to 13.5 mm from the cortical surface. These coordinates corresponded to the dorsal hippocampus, with the vertical tracks passing through *cornu ammonis* (CA) and then the *fascia dentata* (FD).

Although a general trend for functional separation between the dorsal (primarily “cognitive”) and ventral (primarily “visceral”) hippocampus exists in many mammals (Fanselow and Dong, 2010; reviewed by Strange et al., 2014), this transition is smooth. In cats, the dorsal hippocampus is influenced by respiration (e.g., Poe et al., 1996) and its stimulation can induce apnoea. Examination of afferent connections to the cat hippocampus has demonstrated that dorsal portions receive inputs from brain sites associated with viscerosensation, such as the ventroposterior thalamus and the cingulate and orbitofrontal cortices (Irle and Markowitsch, 1982), suggesting that they process interoceptive signals. Although some of the visceral signals, such as breathing can be perceived exteroceptively, especially in wakefulness, for simplicity we refer to visceral-locked hippocampal modulation as interoceptive processing or interoception.

We chose to target the dorsal rather than the ventral hippocampus primarily for easier access via vertical electrode penetrations. This approach was important in our study to limit potential brain damage, which would be more likely to occur with deeper penetrations required for sampling the ventral hippocampus over the study’s duration. It is quite possible that recording activity from the ventral hippocampus would have yielded a higher estimate of the proportions of hippocampal cells synchronized to different visceral events. However, our focus on the dorsal hippocampus provides at least a minimal view of hippocampal synchronization to visceral events.

### IMPLANTATION OF DUODENAL AND GASTRIC ELECTRODES

Bipolar electrodes made of silver were implanted into the walls of the duodenum and stomach (under the serous membrane) to record myoelectrical activity from these organs. The procedure has been previously described by Papasova and Milenov (1965), Papasova et al. (1966), and Pigarev et al. (2013). In short, this laparotomy procedure was conducted under general anaesthesia and aimed to place flattened and polished silver electrodes under the serous membrane of the duodenal wall to access activity of the muscularis externa. The electrodes were secured in place using absorbable sutures penetrating serosa only. These bipolar electrodes had a 2 mm inter-electrode distance, as a shorter distance minimizes signal disturbances caused by postural changes during wakefulness. The electrode leads, made of thin and flexible insulated wires, were passed under the skin using a tunneler to a connector attached to the halo frame. The leads were coiled under the skin to enable posture changes without the leads being pulled from the duodenal or stomach wall. These leads, however, had a limited lifespan due to wear and tear from the animal’s movements and peristalsis as well as the suture absorption time.

To ensure signal quality and duration of recordings necessary to obtain sufficient amount of data, a pair of bipolar electrodes was implanted, 5 cm apart, in the duodenal wall. The best signal was later selected for analysis in one of the animals. In the other animal, both duodenal electrodes provided good-quality signals throughout the entire experimental period, and both signals were used for further analysis. Data collection commenced after a recovery period of at least 4 weeks.

### DATA ACQUISITION

The animals were fed a half of a standard can of wet cat food (approx. 40-45g portion) before the start of each recording. Satiety is known to induce sleepiness and this approach nearly guaranteed that SWS would be recorded during a daily session.

Hippocampal activity was recorded using a pair of tungsten varnish-coated microelectrodes in a bipolar mode. The two electrodes had impedances ranging from 0.5 to 1.5 MΩ and were implanted with a 300 μm distance between their tips. They were inserted together through conical guiding tubes and moved along the vertical axis using a custom-made microdrive attached to the halo frame. Such bipolar recording increases the neuron/recording yield and also allows for the analysis of local field potentials. The electrodes were repositioned prior to every recording session to access a new group of hippocampal neurons. Each repositioning was at least 200 microns but could be larger if no clear spiking activity was observed after the move.

Raw signals were amplified by a bipolar amplifier (NeuroBioLab) in the 0.3–3000 Hz range, sampled at 10,000 Hz, and stored for offline analysis. These neuronal recordings were then filtered offline in the 0.3–250 Hz range to analyse local field potentials (LFPs) and in the 300–3000 Hz range to assess spiking activity. To distinguish action potentials from different hippocampal neurons surrounding the electrode tips, neuronal signals were spike-sorted using the inbuilt algorithm of Spike 2 software (Cambridge Electronic Design Limited), which takes into account spike amplitude, polarity, and waveform shape (for further details and an example of the process, see Levichkina et al., 2021). Spike 2 software has an inbuilt check for ISI violation and provides PCA visualization tool that were used during spike sorting. As discussed by Pedreira et al. (2012), the maximum number of correctly identified neurons by this type of algorithm typically ranges between 6 and 10 cells due to algorithm saturation, although the number of cells in the tissue close to the electrode tip can be much higher. This also varies depending on electrode impedance and tissue geometry. In our experiments, cell yield per recording site varied between 3 and 9 neurons.

Duodenal and gastric myoelectrical signals were amplified in the 0.3 to 70 Hz range and sampled at 200 Hz. To record the ECG, we used one of the duodenal electrodes and the epidurally implanted grounding screw. The signal was sampled at 1 kHz. Signals related to respiration were obtained from the nasal airflow, registered via thermo-sensors placed in front of the animal’s nose, with a 200 Hz sampling rate. The frequency range for both signals was 0.3 to 70 Hz. An infrared oculometer was utilized to record eye movements at 200 Hz sampling rate.

## DATA ANALYSIS

### STATISTICAL ANALYSIS

We used a bootstrapping approach to evaluate the statistical significance of the relationship between neuronal spike trains and visceral events for the analysis of (i) hippocampal neuronal responses triggered by visceral events (ii) visceral activity modulated by hippocampal neuronal spikes and (iii) hippocampal LFP responses to periodic visceral events. Further details of the procedures are provided in the corresponding sub-sections of the Data Analysis section. In short, it included the following steps:

1. Subdividing simultaneous recordings of hippocampal, mainly aperiodic, signals and the visceral, mainly periodic, signals into two sets, one for the SWS intervals and one for wakefulness intervals.
2. For each of the above sets (wakefulness and sleep), calculation of average signal amplitude changes over a period common to the visceral signal in question (say, a respiratory cycle) or the empirically chosen corresponding intervals for the aperiodic signal train.
3. For event-triggered analysis – calculating the number of visceral events that occurred within the wakefulness or sleep interval and placing the same numbers of pseudo-markers over the same intervals for triggering “random” averaging that is event-unrelated. For spike-triggered analysis, the number of pseudo-markers was equal to the number of neuronal action potentials that occurred over the interval in question.
4. Repeating the process 1000 times, 1000 pseudo-modulation values of the signal amplitude were calculated for every cell/visceral signal.
5. The actual amplitude modulation was then classified as significant if it exceeded 95% of the pseudo-modulation values.

To analyse predisposition of the LFP responses to micro-sleep and micro-wakefulness episodes we used Similarity Index (see Fedorov et al., 2023 for a detailed description of the method and the freely available Matlab code, https://github.com/george-fedorov/erp-correlations). High similarity index indicates that an individual (single-trial) LFP has a shape close to the averaged LFP response. With this method, we separated individual LFP responses to visceral events into high-similarity (r²>0.4) and low-similarity groups (r² < 0.1) to further compare the spectral composition of hippocampal activity close to where the LFPs occurred between these two groups using the Wilcoxon rank sum test. The goal of this comparison was to establish whether high-similarity LFPs belong to a particular state of vigilance or distributed randomly between them. Details of this analysis can be found in the sub-section titled “LFP responses to periodic visceral events and vigilance state dependency of LFP responses”. χ2 with Yates correction was applied to test for predispositions of neuronal activity to a particular state of vigilance.

We used the Wilcoxon rank-sum test (p<0.05) to test action potential shapes for electrode displacement, using the approach of Mosher et al. (2020). For details, see “Spike shape test for electrode displacement” section. All analyses were conducted using Matlab software package version R2019B (RRID:SCR_001622).

Our experimental design, based on recording of natural (in contrast to experimentally elicited) signals, did not allow us to solve “chicken or the egg” problem – we could only establish presence or absence of synchronization between visceral and hippocampal signals, not the direction or any common driving force for such synchronization.

### STATE OF VIGILANCE ASSESSMENT

The animals were video-monitored during the recording sessions to assist in estimating the state of vigilance as well as in artefact rejection. The state of vigilance was assessed twice independently, first preliminarily during recording by direct observation of the animal’s behaviour and brain activity to ensure the presence of the necessary states of vigilance, and then off-line using EEG, ECG, and the data on eye movements and breathing pattern. The presence of high-amplitude delta waves in the EEG signal, absence of bodily movements, reduction in amplitude and slowing of eye movements, decrease in heart rate and slow and regular breathing indicated SWS. Open eyes with the presence of saccades, overall alertness (e.g., redirection of the ears towards new sounds), desynchronized EEG and increased frequency and variability of both heart rate and breathing pattern indicated wakefulness. In the analysis described further, we used periods of recording containing either SWS or wakefulness without motion artefacts. Transitional states and REM were excluded from the analysis since due to their shorter durations these states had drastically fewer visceral activity periods available for the analysis.

In cat 1 we recorded between 1 and 4 sleep-wake cycles per recording session (N complete daily sessions = 10), and in cat 2 – from 1 to 7 sleep-wake cycles per recording session (N complete daily sessions = 12). Total length of recording ranged from 1319 to 7367 seconds in cat 1 and from 607 to 4322 seconds in cat 2. Usable recording time per session (after excluding artifacts and transitional states, and checking for data quality allowing placement of the visceral activities markers) ranged from 922 to 3383 sec in cat 1 and from 418 to 2720 sec in cat 2. Usable time in wakefulness ranged from 417 to 1782 sec in cat 1 and from 153 to 846 sec in cat 2, in sleep in was ranged from 177 to 1975 sec in cat 1 and from 503 to 2067 sec in cat 2.

### VISCERAL EVENT-TRIGGERED NEURONAL RESPONSES

For each hippocampal cell, we assessed whether its spike rate was modulated by periodic visceral events, such as heart rate, breathing and duodenal myoelectrical activity. The analysis was performed separately for intervals of sleep and wakefulness within each recording session. Markers for each type of visceral activity were created using a templating process similar to the spike sorting method using the Spike 2 software. The process involved selecting a typical shape of the periodic activity in question and creating a template, with each templated event serving as a marker for further analysis.

We excluded from the analysis those intervals during which the signal changed in such a way that markers could not be reliably constructed. This was necessary because motion artefacts produce events that could be misconstrued as genuine visceral events. Moreover, for duodenal activity, a change in the animal’s posture or a period of vigorous peristaltic activity could sometimes substantially alter the shape of the signal, although its periodic structure remained clear. In such recording sessions, the most consistent shape was used as a template of duodenal activity. Figure 1.C. demonstrates examples of raw visceral signals and the markers derived from these signals.

Duodenal myoelectric activity reflects two physiologically distinct types of signals. The first is a low-frequency activity called simple waves (or electrical control activity), which represents the activity of the enteric nervous system. This type of activity is related to the possibility of duodenal wall contractions and defines their timing, but it occurs regardless of any actual contractions and is intrinsically periodic. We further refer to this type of activity as “simple waves (SW)”. The second type, high-frequency spike bursts (also known as electrical response activity), is a superimposed higher-frequency activity commonly related to the actual contraction of the intestinal wall (Papasova and Milenov, 1965; Papasova et al., 1966; Costa and Furness, 1982; Sarna, 1989; Martinez-de-Juan et al., 2000). This type of activity is referred to as “simple waves with bursts (SB)”. Correspondingly, the first type of activity – SW – is common for the intervals without peristaltic activity, during which slow digestive breakdown of food occurs in the intestinal lumen, while the second one – SB – can be related to intervals when a food pellet is mixed within or moved along the intestine. Thus, different signals can potentially be sent from the intestinal wall to the brain via the vagus nerve when either simple waves or bursting activity dominates.

In the postprandial period, contractions occur at an approximate rate of 4–5 per minute, while the period of simple waves is usually only about 3 seconds long. Typically, a contraction is associated with several episodes of bursting activity, some of which precede the actual contraction. The energy of simple waves is concentrated below 2 Hz, while for the bursts of spike potentials it is above 2 Hz, with most of their energy within the 2–10 Hz range (Sarna, 1989; Martinez-de-Juan et al., 2000; Garcia-Casado et al., 2005, 2006).

After assigning a marker to every simple wave, the resulting markers were separated by spectral analysis into two groups, as in our earlier studies (Levichkina et al., 2006; Pigarev et al., 2013): (1) true simple waves (SW) and (2) waves that also contained higher-frequency bursts (SB). In our experience, the best separation of higher-frequency bursts from simple waves can be achieved by band-pass filtering the raw signal in the 6–35 Hz range and using a sum of the spectral power to reveal the bursts superimposed on simple waves. We then used these two types of duodenal activity markers independently to trigger spike rate averaging of hippocampal activity.

### STATISTICAL ANALYSIS OF THE VISCERAL EVENT-TRIGGERED NEURONAL RESPONSES

The hippocampal spike train was binned every 1 ms, smoothed by a Gaussian kernel with a sigma of 20 ms and averaged using markers of visceral activity as triggers. The highest spike rate difference between two points within one period of the observed visceral signal was taken as the putative value of spike rate modulation. Since the resulting spike rate difference could be caused potentially by a random variation of the spike rate, it was necessary to test the significance of such modulation. To do so, we randomly placed the same number of pseudo-markers for each SWS and wakefulness period in a particular recording session as the number of activity periods derived from the observed periodic visceral signal for the same interval. We then used these pseudo-markers for triggering spike rate averaging. This process was repeated 1000 times for every cell across intervals of sleep and wakefulness. As a result, 1000 pseudo-modulation values were calculated. The actual spike rate modulation over a period of a visceral signal was deemed significant if it exceeded 95% of these pseudo-modulation values.

### SPIKE SHAPE TEST FOR ELECTRODE DISPLACEMENT

Some periodic visceral signals, such as breathing and blood pressure changes associated with heartbeats, may have the potential to displace brain tissue in the vicinity of an electrode due to brain pulsations caused by them. These mechanical changes may, in turn, lead to alterations in the recorded spike shapes (Mosher et al., 2020). As spike sorting algorithms rely on spike shape templating and allow a pre-set amount of noise to be superimposed on a spike shape, these shape changes can potentially translate into algorithm performance variations along the periods of breathing and heart rate, causing artifacts in spike rate estimation. The presence of such changes depends on various factors, including the animal’s brain size, the proximity of an electrode to large blood vessels, and the “stickiness” of an electrode – different electrode shapes and materials may result in varying propensities to either move with the tissue or move away from it. Our bipolar electrode, with two electrodes manufactured to have small balls at their tips and inserted together, was designed to reduce the displacement effect. However, it was still necessary to test whether the observed significant changes in spike rates were associated with the transfer of visceral information or simply reflected these so-called pulsatile artefacts.

To test this, in every trial (a single period of either heart rate or breathing rate), we took all hippocampal spike shapes from the intervals surrounding the identified minimum and maximum of the averaged spike rate along the period of the signal in question. The interval was equal to 20 ms (±10ms from a peak or a trough) in most cases. However, for the rarely spiking cells, we increased it to 30 ms to collect enough spikes for further comparisons. Mosher et al. (2020) analysed multiple parameters of the extracellular spike shape and demonstrated that the two most sensitive to tissue displacement ones are the spike amplitude (AMP) and its half-width (HW). We calculated these parameters as suggested by Mosher et al. and compared these values between the spikes surrounding a peak and those surrounding a trough using the Wilcoxon Rank Sum test. If the spike shapes in our case were indeed distorted by tissue displacement, the highest spike shape difference was expected to be between these intervals. This analysis was performed individually for each of the cells that demonstrated significant changes in their spike rates along either the heart or breathing periods, but not for the duodenum rate-related cells since duodenal peristaltic activity cannot displace brain tissue. The analysis revealed only a minor presence of the displacement effect in our data: We found at least one of the two spike shape parameters to be significantly different (p<0.05) for 4 cells for the breathing-associated spike rates (3 of these 4 cells belonged to the same recording site) and 2 for the heart rate. These cells were excluded from the results presented in this paper.

### SPIKE-TRIGGERED VISCERAL ACTIVITY MODULATION

The activity of hippocampal neurons might be associated with specific visceral events, such as a deep sigh, rather than aligned with the period of regular visceral events like breathing rate. Additionally, some visceral signals, such as stomach myoelectric activity, may lack stable periodicity. To investigate the relationships between hippocampal cell activity and these types of visceral events, we performed a spike-triggered average (STA) of visceral activity. This analysis was conducted for stomach myoelectric signals, breathing, averaged heart rate, and the envelope of the high-frequency component of the duodenal signal, which is related to the preparation and execution of peristalsis. Spike triggering can reveal whether spiking changes are aligned with changes in the amplitude of a visceral signal rather than its period.

STA was computed using the timing of every spike from a particular neuron as a trigger for averaging the visceral signal. For periodic signals of varying amplitudes, the goal of STA analysis was to test for a possible link between high-amplitude visceral events and neuronal spiking frequency. In this case STA amplitude was measured at an interval equal to a period of a visceral signal in question +1 second to account for a possible variability of a period. For aperiodic signals such as stomach myoelectric activity and mean heart rate, the interval of averaging was chosen arbitrarily based on the observed natural intervals of modulation for these signals. We chose ±6 second period for stomach activity and ±30 seconds for heart rate.

The flanking parts of the vigilance state interval of the analysis were excluded from STA calculation to prevent mixing vigilance states during averaging. These excluded parts were equal to half of the period of averaging at the beginning and end of the vigilance state interval.

The envelope of the high-frequency component of the duodenal signal was calculated by detrending the raw signal, bandpass-filtering it (6 to 35 Hz, Butterworth, 6th order, bidirectional), and taking the absolute value of a Hilbert-transformed signal.

Heart rate was calculated as a sum of heartbeats occurring in 10 seconds, with a 10ms step. Stomach activity could be contaminated by breathing when diaphragm movements affect the area of the electrode placement. That effect depended mainly on the animal’s posture. If a hippocampal neuron actually responds to a breathing rate or to sighing, the STA of stomach activity would show the breathing dependency rather than reflect stomach activity. Although stomach activity could have frequency components similar to the breathing period, it was impossible in our experiment to distinguish the origin of STA in such cases. Therefore, we decided to exclude from the stomach activity analysis all STA results where the resulting STA wave had frequencies less than 2 breathing periods. That caused a reduction in the reported stomach-related STAs.

### STATISTICAL ANALYSIS OF SPIKE-TRIGGERED VISCERAL ACTIVITY MODULATION

Statistical analysis was based on bootstrapping. The number of neuronal spikes within any interval in question (corresponding to wakefulness or SWS) was calculated and the same number of random pseudo-spike markers was generated across the same interval and used to calculate pseudo-STA. This process was repeated 1000 times for every cell and 1000 pseudo-STA values were obtained. The actual STA value was considered significant if it exceeded 95% of the pseudo-STA values.

### LFP RESPONSES TO PERIODIC VISCERAL EVENTS AND VIGILANCE STATE DEPENDENCY OF LFP RESPONSES

We also analysed evoked LFPs in a manner similar to the previously described analysis of visceral event-triggered neuronal spiking responses. While neuronal spiking reflects the output of a specific brain region, local field potentials (LFPs) are more closely related to its input, as they represent the summation of dendrosomatic synaptic signals (Logothetis, 2002; Raichle and Mintun, 2006).

However, as LFPs are known to be affected by pulsatile artifacts (Fee, 2000; Mosher et al., 2020), it is usually recommended to filter the LFP signal at a frequency cut-off above the heart rate frequency. The actual metabolic changes associated with brain tissue perfusion and changes in CO2 levels resulting from neuronal activity are also very likely to produce LFP changes, and recent studies have uncovered a mechanism of direct pulse wave detection by mitral cells of the olfactory bulb (Salameh et al., 2024). However, these real changes are difficult to distinguish from pulsatile artifacts in experiments conducted with non-anesthetized animals. Therefore, we used a 5 Hz to 125 Hz bandpass filter (Butterworth, 6th order, bidirectional) before calculating heartbeat-evoked LFPs. At the same time, it has been established that breathing-evoked LFPs exist without tissue displacement (e.g., reviewed by Tort et al., 2018), and these responses aren’t normally regarded as artefact-produced. Thus, we used a 0.3 to 125 Hz range for the analysis of breathing-related as well as duodenal activity-related LFP responses to avoid losing low-frequency components of the evoked potentials. Bootstrapping-based analysis was conducted separately for sleep and wakefulness intervals, as described above.

### STATISTICAL ANALYSIS OF THE VIGILANCE STATE DEPENDENCY OF LFP RESPONSES

When significant LFP responses were observed in both vigilance states, we performed additional analysis to determine whether LFP responses had similar characteristic (shapes and amplitudes) across the entire length of the vigilance state in question or occurred during putative micro-sleep or micro-wakefulness episodes. Instead of pre-defining such episodes, which can be tricky in cats and can only be made by arbitrary thresholding EEG or LFP, we decided to test whether difference in the evoked responses statistically significantly corresponded to differences in LFP spectra. We refer to the episodes of heightened low frequency part of the spectra during wakefulness as micro-sleep episode and the opposite – diminished low frequency activity during SWS without transitioning to REM – as micro-wakefulness episodes.

For this analysis we sorted individual LFP shapes trial-by-trial into groups of low and high similarity with the averaged evoked LFP response using an r²-based Similarity Index (see Fedorov et al., 2023 for a detailed description of the method and the freely available Matlab code). We compared the spectra of high-similarity trial LFPs (r²>0.4) to the spectra of low-similarity trial LFPs (r²<0.1) in two frequency ranges related to sleep: delta (1-4 Hz) and spindles (7-15 Hz), using the Wilcoxon rank sum test. This procedure allowed us to determine whether the trials contributing to the averaged evoked LFP occurred during more drowsy or more alert states.

### NUMBER OF CELLS AVAILABLE FOR THE ANALYSIS

Neuronal activity was recorded from 213 individual hippocampal cells (135 cells from cat 1 and 78 cells from cat 2). As described above, STA of stomach activity could only be analysed if it was not contaminated by breathing, so we restricted this part of the analysis to 107 cells. Additionally, we limited the analysis of breathing-related neuronal activity to 184 cells due to poor breathing signals in 2 recording sessions and the exclusion of 4 cells with spike shapes influenced by tissue displacement. For the remaining types of recorded visceral signals, relationships between visceral and hippocampal activities were analysed for each of the 213 cells.

Both duodenal electrodes were preserved in cat 2 for the entire duration of the experiment. We studied the relationships between duodenal and hippocampal activities using signals from both electrodes since different sections of the duodenum do not act synchronously.

We presented the results as percentages of cells that had their activity significantly modulated along a period of the visceral activity in question or as a percentage of cells that triggered significant visceral STAs (above 95% confidence interval, based on the bootstrapping analysis) and as proportions of cells having such significant modulation. Assuming the 5% error “built-in” to the statistical method (p=0.05), one can expect 5% of cell modulations to be deemed significant despite actually being random. Thus, reporting the results as percentages allows the reader to get a correct impression of the results’ validity. Additionally, we report percentages of cells with modulation significance >1% (p<0.01) to further clarify the results.

### CORRECTION FOR MULTIPLE COMPARISONS

The problem of multiple comparisons was tackled in two steps. First, binomial enrichment test was used for each one of the bootstrapping-based results reported (e.g., separately for cells modulated by breathing in SWS vs modulated by breathing in wakefulness, etc.). Enrichment test allows to estimate whether a category of outcomes occurs more frequently than expected by chance in a particular dataset (e.g., Weinstein et al., 2024; Li et al., 2024; Ferguson et al., 2025). The resulting binomial enrichment test p-values were further corrected for multiple comparisons using the Benjamini–Hochberg FDR procedure across all binomial tests performed (q < 0.05 considered significant).

## RESULTS

Basal frequency rates of the registered visceral activities for each of the cats are presented in Table 1, separately for SWS and wakefulness. HR and BR rates were within ranges commonly reported for domestic cats, SW data for various parts of the intestinal tract in cats (e.g. specifically for the duodenum) are too rare in the existing literature to provide a standard background.

**Table 1.** Basal frequencies of the registered rhythmical interoceptive signals.

|  | SWS |  | Wakefulness |  |
| --- | --- | --- | --- | --- |
|  | Cat 1 | Cat 2 | Cat 1 | Cat 2 |
| BR, Hz | 0.35 | 0.46 | 0.43 | 0.52 |
| HR, Hz | 2.12 | 2.72 | 2.29 | 2.82 |
| SW, Hz | 0.31 | 0.45 | 0.31 | 0.45 |

Mean breathing rates were significantly higher during wakefulness comparing to SWS in each of the animals (Wilcoxon signed-rank test, cat 1 p= 0.0117, mean BR in wakefulness 0.43 Hz, BR in SWS 0.35 Hz; cat 2 p = 0.002, BR in wakefulness was 0.52 Hz, in SWS 0.46 Hz). Similarly, heart rates changed between wakefulness and sleep in both animals as well (Wilcoxon signed-rank test, cat 1 p= 0.0195, mean HR in wakefulness 2.29 Hz, HR in SWS 2.12 Hz; cat 2 p = 0.0049, HR in wakefulness 2.82 Hz in SWS 2.72 Hz). These results further confirm the quality of state of vigilance classification. The differences in absolute values between animals were due to the disparity in their corresponding body sizes – cat 1 had larger body frame and body weight. Duodenal baseline rate (SW) did not change significantly between states of vigilance (p>0.05 for each of the animals). SW rate in cat 1 was 0.31Hz in both states, while in cat 2 it was 0.45 Hz in both states as well.

Gastric rhythmicity is intrinsically slow (approx. 3 per minute in humans and 4 per minute in cats, Xue et al., 1995) with amplitude, velocity, and frequency fluctuating due to neural/hormonal inputs, distension, and regional gradients, making it too difficult to use for event-triggering analysis, especially since intervals of deep SWS and clear wakefulness in cats often last only a few minutes. We therefore used only spike-triggered approach to analyse stomach activity.

Mean background spike rates for all the recorded hippocampal cells were 20.6 (SD±21.4) in wakefulness and 15.9 (SD±19.2) in SWS. This difference in spiking was significant (p=0.012, Wilcoxon rank sum test). Previous cat studies demonstrated vigilance-state modulation of hippocampal spiking, with reduced pyramidal discharge in deep SWS in some recordings from dorsal hippocampus and the opposite effect – strong SWS-linked spike rate enhancement in ventral hippocampus (Noda et al., 1969; Hartse et al., 1979; Marczynski et al., 1980). Our results are consistent with previously estimated ones from feline dorsal hippocampus.

### RESPIRATION-RELATED NEURONAL ACTIVITY

Our results revealed that respiration provided the most significant influence on hippocampal activity. This was not surprising, as the effects of breathing on hippocampal activity was described in various species (Duffin and Hockman, 1972; Frysinger et al., 1989; Poe et al., 1996; Radna and MacLean, 1981; Herrero et al., 2018; Tort et al., 2018; Folschweiller and Sauer, 2021; Nokia and Penttonen, 2022), and the high magnitude of this effect noted. However, a new effect emerged when we analysed these dependencies separately in wakefulness and sleep. The majority of cells exhibited significant relationships to breathing either in wakefulness or in sleep, but not in both states. The event-triggered analysis showed that 20.1% (37/184) of cells were significantly synchronized to breathing in wakefulness, 22.8% (42/184) in SWS and 3.8% (7/184) in both states. Additionally, 17.9% of them had high modulation significance (p<0.01). These proportions of synchronisation between hippocampal activity and BR were significant according to binomial enrichment test and survived FDR at q<0.001.

Modulation of baseline spiking by a rhythmical event was calculated for every cell as the following: Modulation, % = ((max spike rate – min spike rate) / mean spike rate) * 100) – 100. The modulation value therefore demonstrates the change between maximal and minimal spike rate of a cell along the cycle of a particular visceral activity as a percentage of its mean rate. We conducted these calculations separately for each state. Mean modulation for cells with significant synchronisation to BR in wakefulness was equal to 25% (SD±36), while in SWS it was 17% (SD±14.8).

Figures 2 and 3 show examples of significant relationships between cell spike rates and breathing. For clarity, all panels in each figure represent the same cell. The cell represented in Figure 2 exhibited strong tonic synchronization to normal breathing in SWS, while the cell demonstrated in Figure 3 responded only to high-amplitude breathing events. Panel 2.A illustrates the animal’s breathing and neuronal spiking of one cell in two states – wakefulness (red) and sleep (blue). The upper part of panel 2.B shows the cell’s averaged activity in wakefulness (red line) and SWS (blue line), while the bottom part of that panel displays the corresponding averaged breathing (registered by the nasal thermistor). The cell’s activity did not appear to be related to breathing during wakefulness, while in sleep, its activity became more regular with patches of spikes aligned with breathing. However, since some cells switch to a “burst-pause” mode in SWS, it was essential to statistically test whether the modulation of cell activity indeed occurs in line with the breathing cycle. This cell indeed had significant breathing-related responses in SWS (p<0.01).

**Figure 2.**
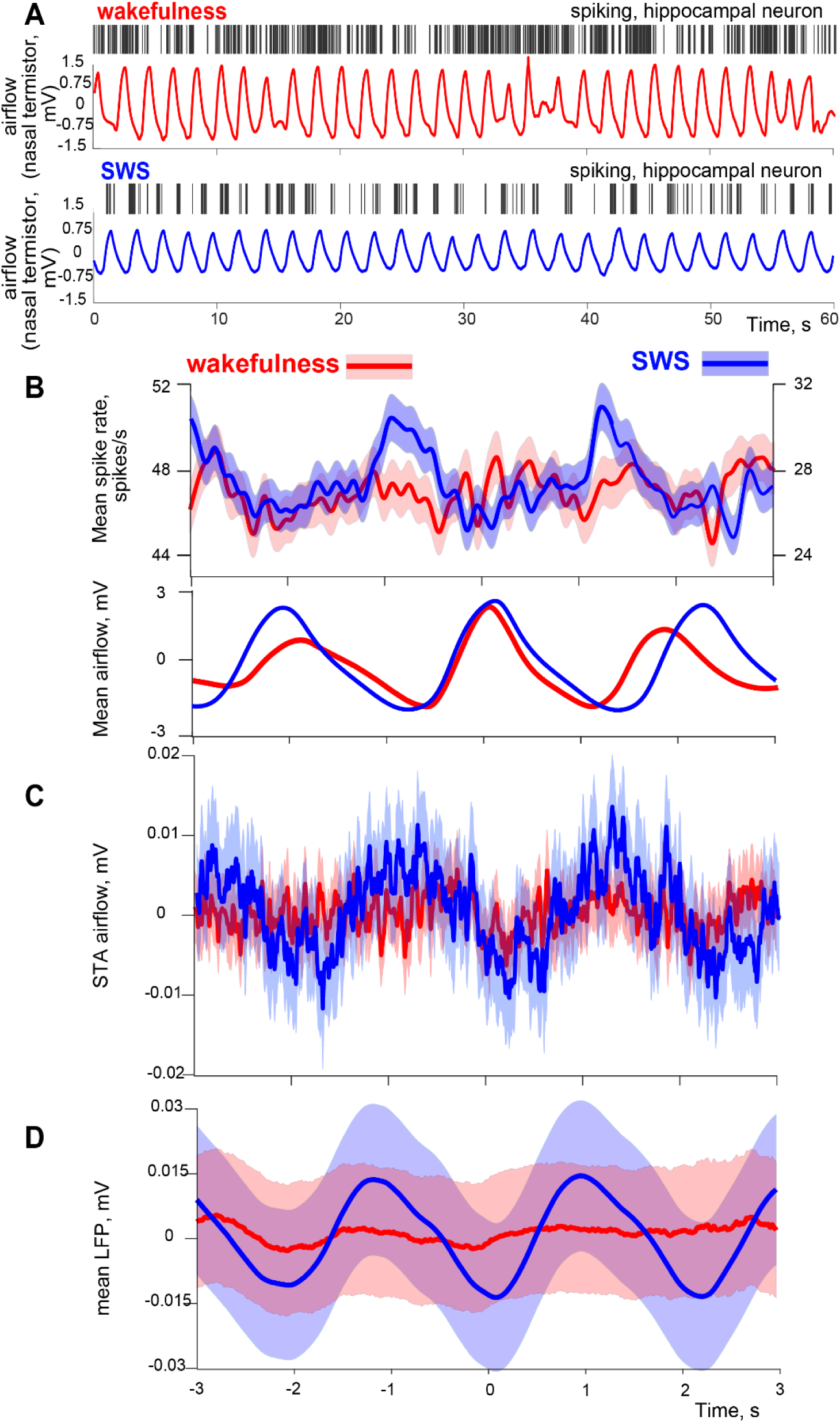
Example of hippocampal activity tonically synchronized to breathing in sleep. A. Breathing and neuronal spiking of a single cell one cell in two states. Air flow measured by nasal thermistor is shown in red during wakefulness in blue in SWS. Individual cell action potentials are represented by vertical lines above the corresponding breathing signal. B. Top panel: event-triggered activity of the same cell, air flow markers served for event-triggering. Solid lines show spike rate average; transparent areas represent the standard error. Bottom panel shows average air flow signal. Blue is reserved for sleep and red for wakefulness. These colours and transparency are used in the same way throughout the Results figures. C. STA of the air flow signal triggered by activity of the same cell. D. Event-triggered response of hippocampal LFP to breathing. The LFP was derived from the same raw signal as the cell activity depicted above (same recording site).

**Figure 3.**
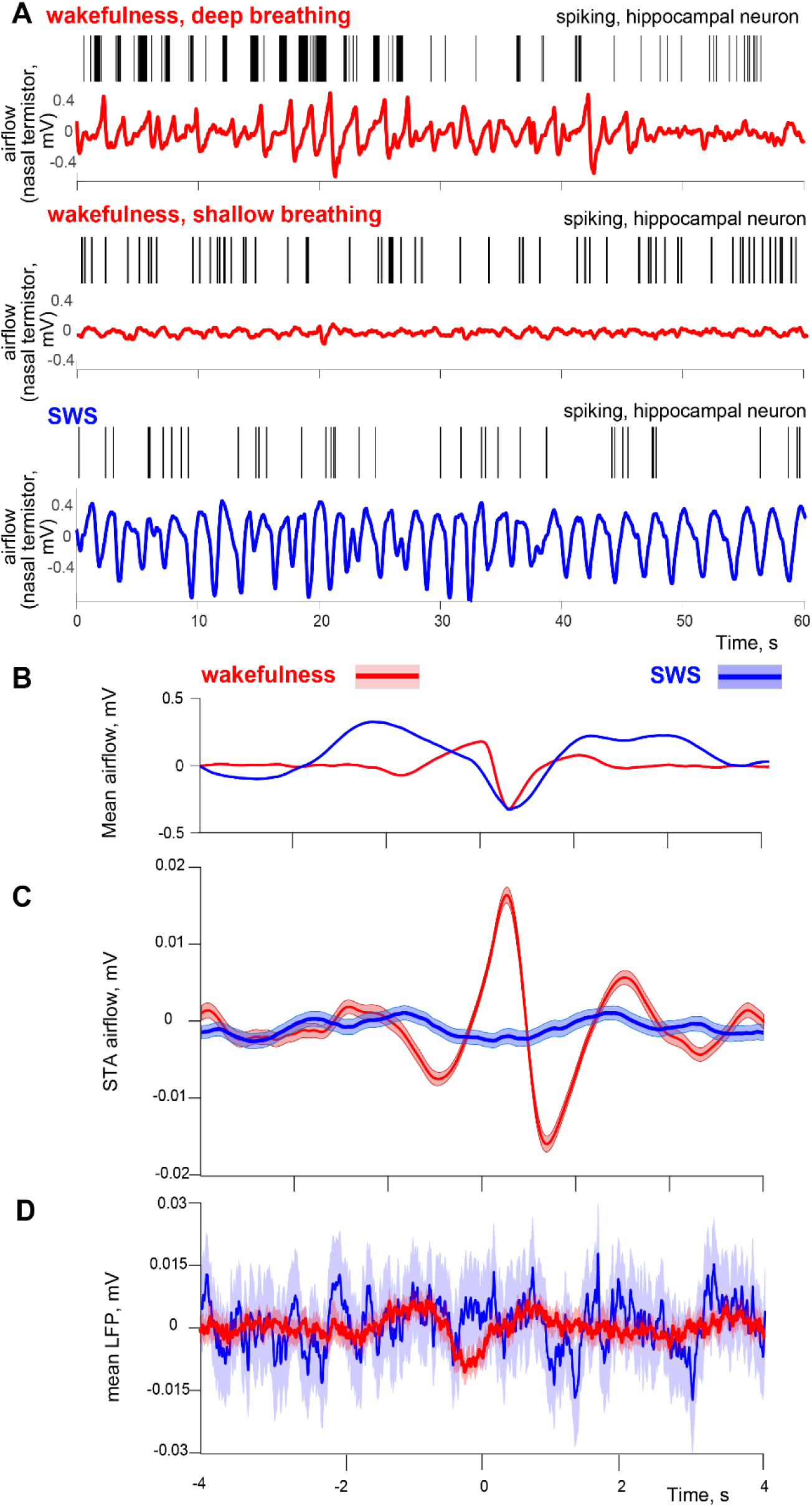
Example of hippocampal activity selectively synchronized to high-amplitude breathing in wakefulness. A. Breathing and neuronal spiking of a single cell in SWS and wakefulness. Top panel – deep breathing in wakefulness, middle panel – shallow breathing in wakefulness, bottom panel – deep breathing in SWS. Air flow measured by nasal thermistor is shown in red during wakefulness in blue in SWS. Individual cell action potentials are represented by vertical lines above the corresponding breathing signal. B. Mean air flow signal. Note that its average amplitude is not higher in wakefulness. C. STA of the air flow signal triggered by activity of the same cell. Event-triggered response of hippocampal LFP to breathing. The LFP was derived from the same raw signal as the cell activity depicted above (same recording site).

Activity of the hippocampal cells could be related to some specific events rather than following breathing activity regularly. While the cell represented by Figure 2.B was tracking breathing in a tonic fashion in SWS, its STA (Fig 2.C) demonstrates moderate, although significant, modulation in SWS as well. In contrast, Figure 3.C provides a dramatic example of such a relationship for a cell that responded specifically to high-amplitude respiratory events and only during wakefulness. To highlight this dependency, panel 3.A shows the cell’s spiking and 3 corresponding episodes of breathing: high amplitude in wakefulness, high amplitude in SWS, and shallow breathing in wakefulness. One can see that only high-amplitude respiratory events during wakefulness were associated with a substantial increase in the cell’s spike rate. The cell’s spiking rate dramatically increased when breathing amplitude was high during wakefulness, but not in SWS.

STA analysis revealed a larger difference between states than the event-triggered one, with 23% (42/184) of cells in wakefulness, 42.9% (79/184) in SWS and 12% (22/184) in both. 45.2% of them had high modulation significance (p<0.01). These proportions of synchronisation between hippocampal activity and breathing were significant according to binomial enrichment test and survived FDR at q<0.001 just like for the event-triggered analysis.

This sleep preponderance of STAs was highly significant (χ2 with Yates correction = 7.38, p = 0.006) for the entire population of cells as well as for each of the animals individually (cat 1 χ² = 8.7, p = 0.003; cat 2 χ² = 5.64, p = 0.017).

A high proportion of cells (21.8%) demonstrated significant synchronization to regular breathing in conjunction with significant STA in the same vigilance state. This indicates that the level of cell spiking modulation by breathing can change over time, even within the same vigilance state.

We also observed significant synchronization of LFPs to breathing (40% of recording sites in SWS and 47% in wakefulness, surviving enrichment test and FDR at q<0.001). However, unlike individual cells, LFPs were often synchronized to breathing in both vigilance states, with 23% of recording sites showing significant synchronization in both states. This prompted further investigation into the preference of LFP responses to micro-sleep or micro-wakefulness episodes.

The analysis revealed a preponderance of LFPs with significant relationships to breathing to episodes of higher delta power (Wilcoxon rank sum test, p<0.05). The majority of such recording sites tended to have pronounced LFP responses to breathing during deeper sleep states in SWS and higher levels of delta activity during wakefulness, likely representing micro-sleep/drowsiness states punctuating wakefulness. Among the 23% of recording sites which synchronized to breathing in both states, 20% showed synchronization during higher delta activity in SWS and 17% during higher delta activity in wakefulness. Only one recording site demonstrated the opposite effect during wakefulness, and none during SWS. This result further confirms the strengthened link between breathing and hippocampal activity during sleep and drowsiness. Examples of LFP responses to breathing in sleep and wakefulness are demonstrated in Figures 2.D and 3.D for the same recording sessions as the other panels of these figures.

### CARDIAC-RELATED NEURONAL ACTIVITY

A moderate number of hippocampal neurons had spiking modulated along the cardiac cycle, and these cells synchronized their activities to heartbeats only in one condition. The event-triggered analysis revealed that 7.1% (15/213) of cells were linked to heartbeats in wakefulness, 8% (17/213) in SWS, and no cells had this relationship in both states. Of these cells, 5.2% had modulation significance >1%. However, these percentages did not meet the thresholds for FDR (q>0.05). Thus, we cannot reliably confirm synchronization between hippocampal spiking activity and individual heartbeats. However, as LFP responses demonstrated high sensitivity to heart rate (see below), it seems likely that some individual cells could also be synchronized to heartbeat, perhaps without maintaining high degree of synchronization all the time but only synchronising to it under some conditions.

Cells synchronized with heartbeats had modulation values in wakefulness equal to –0.4 % (SD± 23.2), in SWS it was 10.9 % (SD±37). Cells synchronising to this activity had the smallest modulation values among all groups of cells.

Figure 4 provides examples of cell activities synchronized to the heart rate cycle in either wakefulness or sleep. Figure 5 demonstrates LFP synchronization to heartbeats in wakefulness and slow wave sleep.

**Figure 4.**
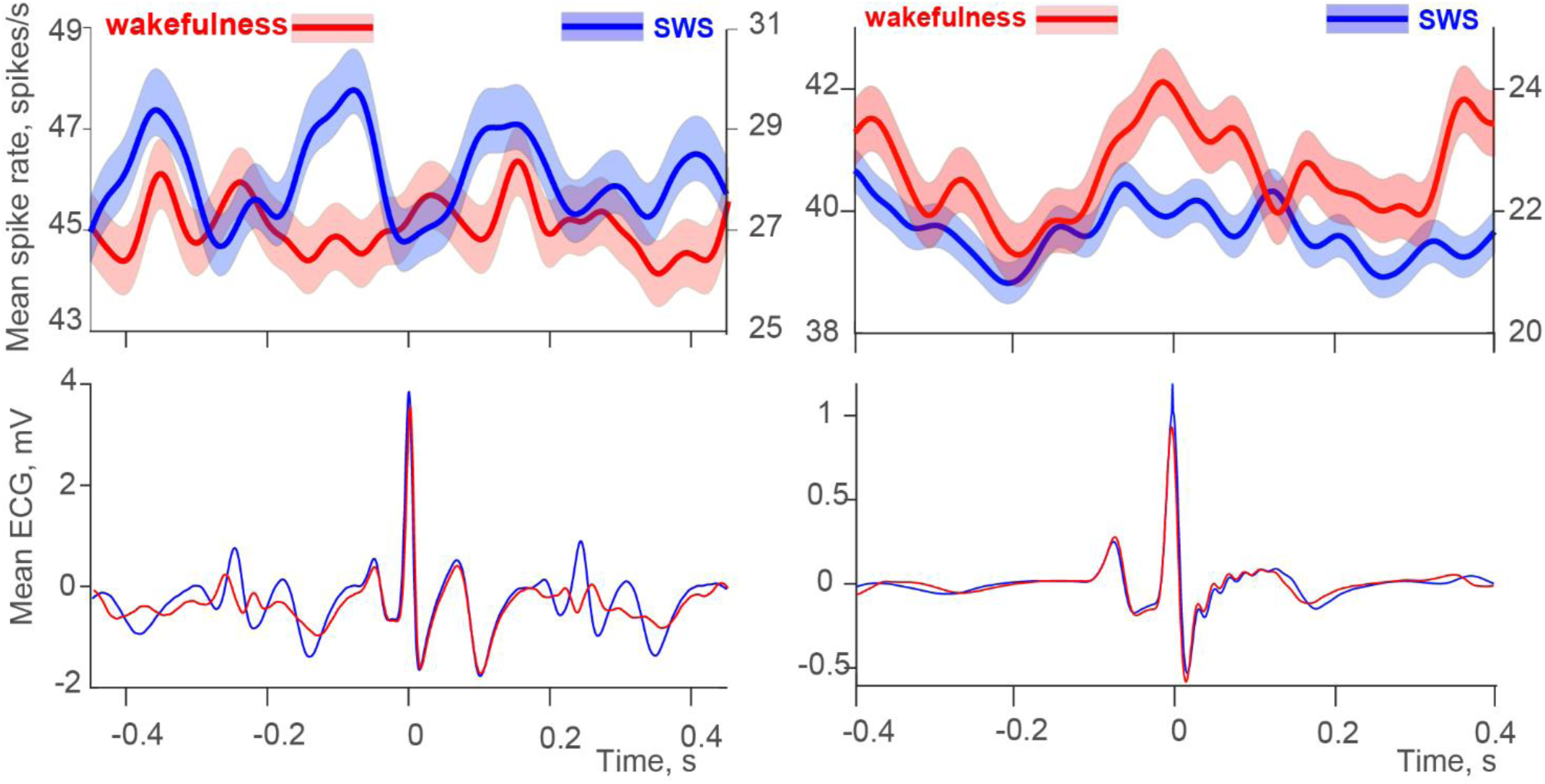
Activities of two hippocampal cells (top row), one synchronized with heart rate cycles in sleep (left side) and the other in wakefulness (right side). The corresponding averaged ECG signals are in the bottom row.

**Figure 5.**
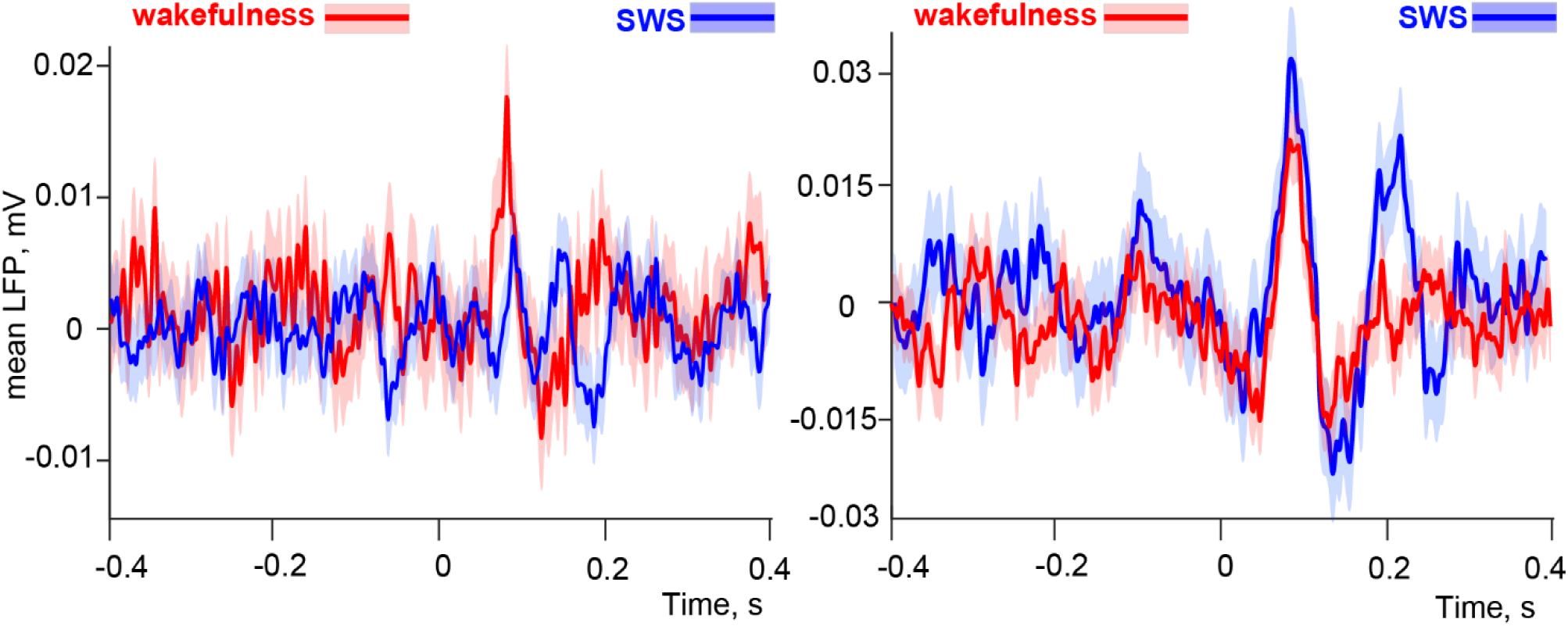
Event-related hippocampal LFPs responses to heart rate cycle, significant in either wakefulness (left side) or sleep (right side).

### JOINT RESPONSES TO BREATHING AND CARDIAC CYCLES

A few cells had relationships to both the cardiac cycle and the breathing cycle within the same states of the sleep-wake cycle. For example, the cell that we show in Fig. 2 as being modulated by breathing in sleep also had its activity synchronised to the heartbeats, as shown by the same cell in Fig 4. Modulation of spiking by two parameters at the same time provides grounds for complex entrainment of the hippocampal cells and potentially being triggered at frequencies that may differ from the ones in the visceral periodic activities.

### STA OF HEART RATE

In contrast to only moderate association between cell spiking and cardiac cycle, STA of the heart rate yielded an unusually high proportion of significant results: 71.8% (153/213) of cells in wakefulness, 70% (149/213) in SWS, and 50.7% (108/213) in both states. However, it is important to note that one cannot draw clear conclusions regarding the relationship between cell spiking and heart rate due to the following two confounds:

(i) Heart rate changes between wakefulness and sleep in a stereotypical way, it is slower and more regular in SWS. Simultaneously, some cells also have state-dependent spike rate modulations, thus in some cases, the changes would co-occur even without a direct relationship between the cardiac cycle and hippocampal activity.
(ii) The hippocampus is involved in emotion regulation, while heart rate is also dependent on the animal’s (or person’s) emotional state. This also provides grounds for co-occurrence of the two rate changes rather than a direct relationship between them.

Nevertheless, a direct relationship is also possible – after all, the hippocampus is well-positioned to affect heart rate due to its emotion-regulation properties and connections with the hypothalamus and other structures involved in emotional and cardiac regulation (Kishi et al., 2000; Petrovich et al., 2001; Herman et al., 2005; Castle et al., 2005; Dong and Swanson, 2006; Fanselow and Dong, 2011). Our experimental setup did not allow us to investigate this question further to distinguish co-occurrences from true relationships. We report this effect mainly to illustrate the prevalence of that co-occurrence, which demonstrates that hippocampal spike rate is highly associated with heart rate frequency, regardless of the driving force behind that association. This can potentially explain the known comorbidities between TLE and emotional disturbances on one hand, and, on the other hand, the well-known predictive value of heart rate in vagus nerve stimulation (VNS) treatment (Jeppesen et al., 2010; Liu et al., 2018; Karoly et al., 2021).

Similar to the LFP effects observed for breathing, LFP responses to heartbeats were significant in 43% of sites in SWS, 40% in wakefulness, and 23% in both states, with both SWS and wakefulness-related proportions surviving enrichment test and FDR at q<0.001 (Figure 5). Thus, their predisposition to microsleep/wakefulness was analysed as well. Among the dual-responding sites, only 7% in sleep and 13% in wakefulness occurred predominantly during episodes of higher delta power. However, in addition, 13% in SWS and 10% in wakefulness preferred episodes of higher power in the range of sleep spindles (7-15Hz), possibly indicating a preponderance to relatively light sleep or transitional states, as spindles are associated with NREM-REM transition in both cats and humans (Gottesmann, 1984; Carrera-Cañas et al., 2019). Thus, in total, nearly half of LFP responses had high preponderance of either delta activity or spindles in both SWS and wakefulness. Note that the rest of the synchronisations with heart beats did not have a trend towards wakefulness (low delta or spindles states) but simply did not demonstrate any significant preferences. LFP synchronization to cardiac events was, to an extent, sensitive to the state of vigilance, demonstrating a preference for sleep similar to the LFP synchronization to breathing, but possibly without a strong preference for deep sleep.

### GASTROINTESTINAL INFLUENCES

#### 1. Duodenum

Duodenal myoelectric signals representing simple waves (SW) and waves with superimposed higher frequency bursts (SB) were marked separately, and hippocampal neuronal activity was subsequently averaged using them as distinct triggers. For one of the two cats, myoelectric activity was recorded from two electrodes, and neuronal activity was analysed using triggering signals from both. The resulting number of analysed pairs of “myoelectric activity – neuronal activity” was 288, thus we reported percentages where 288 is equal to 100%. SWs correspond to periods without peristaltic activity in a particular section of the duodenum, while SBs are to some extent correlated with peristalsis. Consequently, different types of information about the chemical composition of the luminal content and mechanical distention of the duodenal wall should be transmitted to the central nervous system during these periods. The separation between these types of signalling was revealed in our data: only 2% of cells had significant synchronization to both SW and SB simultaneously, and in wakefulness, that proportion was even smaller (1%).

##### 1.1 SB and the envelope of high frequency duodenal activity

Nearly equal numbers of cells synchronized their activities to SB in wakefulness and sleep, again with a very small proportion in both conditions. The percentages were: 10.8% (23/213) in wakefulness, 10.3% (22/213), in SWS and 0.95% (2/213) in both. 2.8% of cells had high modulation significance (p<0.01). Both wakefulness and SWS proportions were significant according to binomial enrichment test, but only wakefulness-related relationship survived FDR (q = 0.033), while SWS FDR was marginal (q = 0.07). Cells synchronized with duodenal SB had modulation values equal to 31% (SD±16) in wakefulness, in SWS it was 42% (SD±38). Figure 6.A demonstrates synchronization between a hippocampal neuron and regular SB signal occurring in sleep. Panel B shows the synchronization occurring in wakefulness for a different hippocampal neuron. LFP responses to SB events were observed in 10% of recording sites in SWS and 12.5% in wakefulness, with 0% in both states. Only the wakefulness-related LFP responses were significant according to binomial enrichment test, and marginal after FDR (q = 0.056). Fig 6.C shows SB-evoked LFP examples.

**Figure 6.**
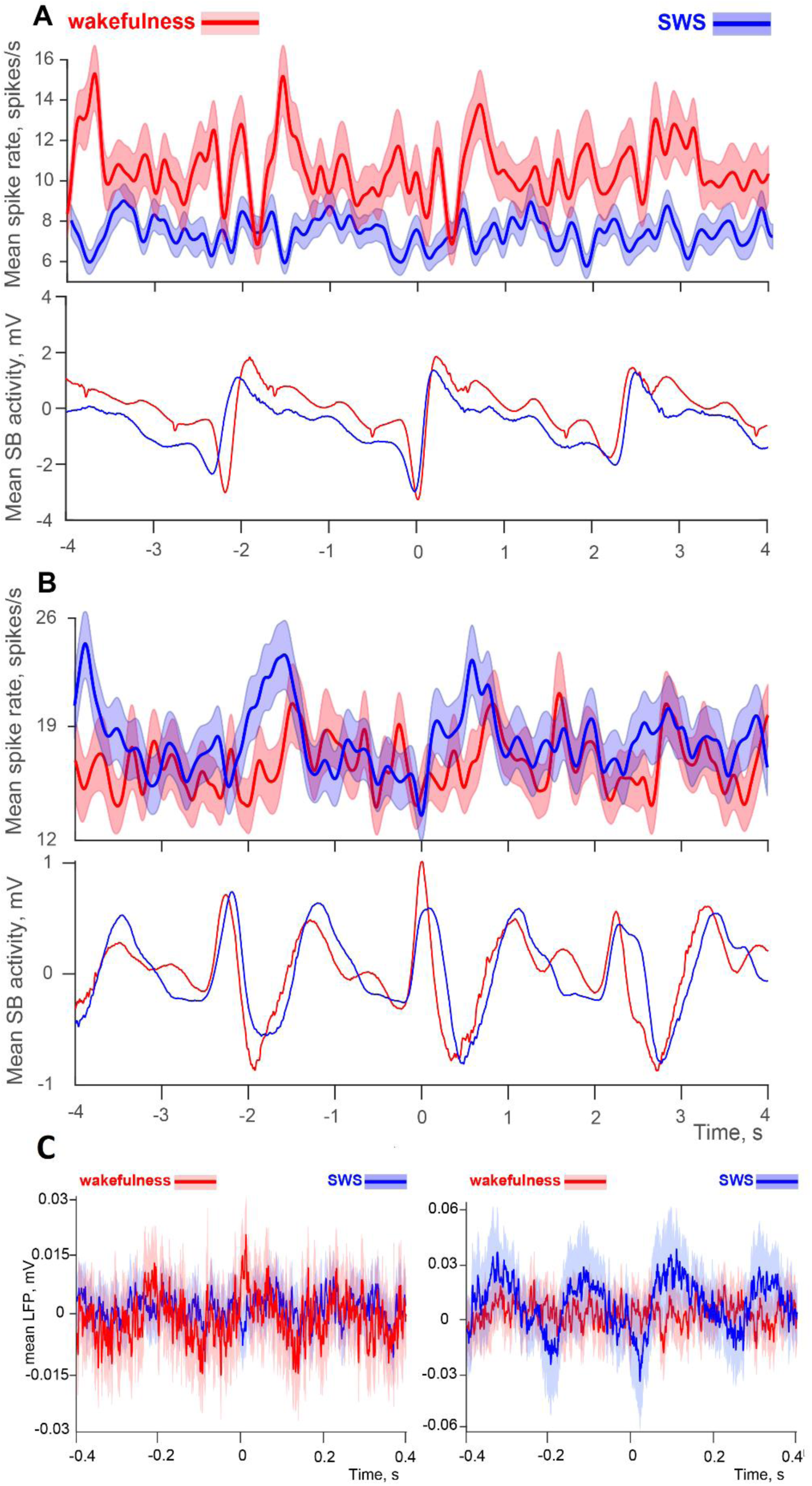
Synchronization of hippocampal activity with SB duodenal signals. A. Activity of a hippocampal cell synchronized to SB in wakefulness, top row. The corresponding averaged duodenal myoelectric activity signals are in the bottom row. B. Activity of a hippocampal cell synchronized to SB in sleep, top row. The corresponding averaged duodenal myoelectric activity signals are below. C. Event-related hippocampal LFPs responses synchronized with duodenal cycle in either wakefulness (left side) or sleep (right side).

For the pooled data significant STA of the SB envelope was more frequently observed in wakefulness (χ2 with Yates correction = 4.24, p = 0.039, 11.3% (24/213) in wakefulness, 6.15% (13/213) in SWS and 1.9% (4/213) in both. 7.3% of them had high modulation significance (p<0.01). In addition, only the wakefulness-related proportion survived both enrichment test and FDR (q = 0.009) while proportion for SWS was not significant (q = 0.338). Due to relatively small amounts of cells demonstrating significant STA in each of the animals the individual χ2 did not reach significance, however, both animals had similar trends towards having more robust STA results in wakefulness. It is expected that not every individual SB can lead to a peristaltic movement, and higher SB amplitudes indicate a higher probability of peristaltic activity. Our results showed a possible connection between peristaltic activity and hippocampal activity occurring in wakefulness. This preponderance to the awake state corresponds to the results showing increased duodenal motility in wakefulness compared to SWS (reviewed by, e.g., Duboc et al., 2020).

##### 1.2 SW

Significantly more hippocampal cells synchronized to SW in sleep compared to wakefulness (χ2 with Yates correction = 8.6, p = 0.003), with 6.6% (14/213) in wakefulness, 15.5% (33/213) in SWS and 0.95% (2/213) in both. 4.2% of them had high modulation significance (p<0.01). SWS-derived proportion was significant after binomial enrichment test and FDR (q = 0.0005) while wakefulness-derived one was not significant at the stage of the enrichment test (p = 0.123). Results for each of two animals were similar: cat 1 χ² = 4.18, p = 0.04; cat 2 χ² = 4.61, p = 0.03.

Cells that synchronized with duodenal SW had modulation values in wakefulness equal to 37% (SD±32), in SWS it was 31% (SD±19). Figure 7 (panels A and B) shows examples of mean spike rates of hippocampal cells selectively synchronizing their activity to SW in either wakefulness or sleep.

**Figure 7.**
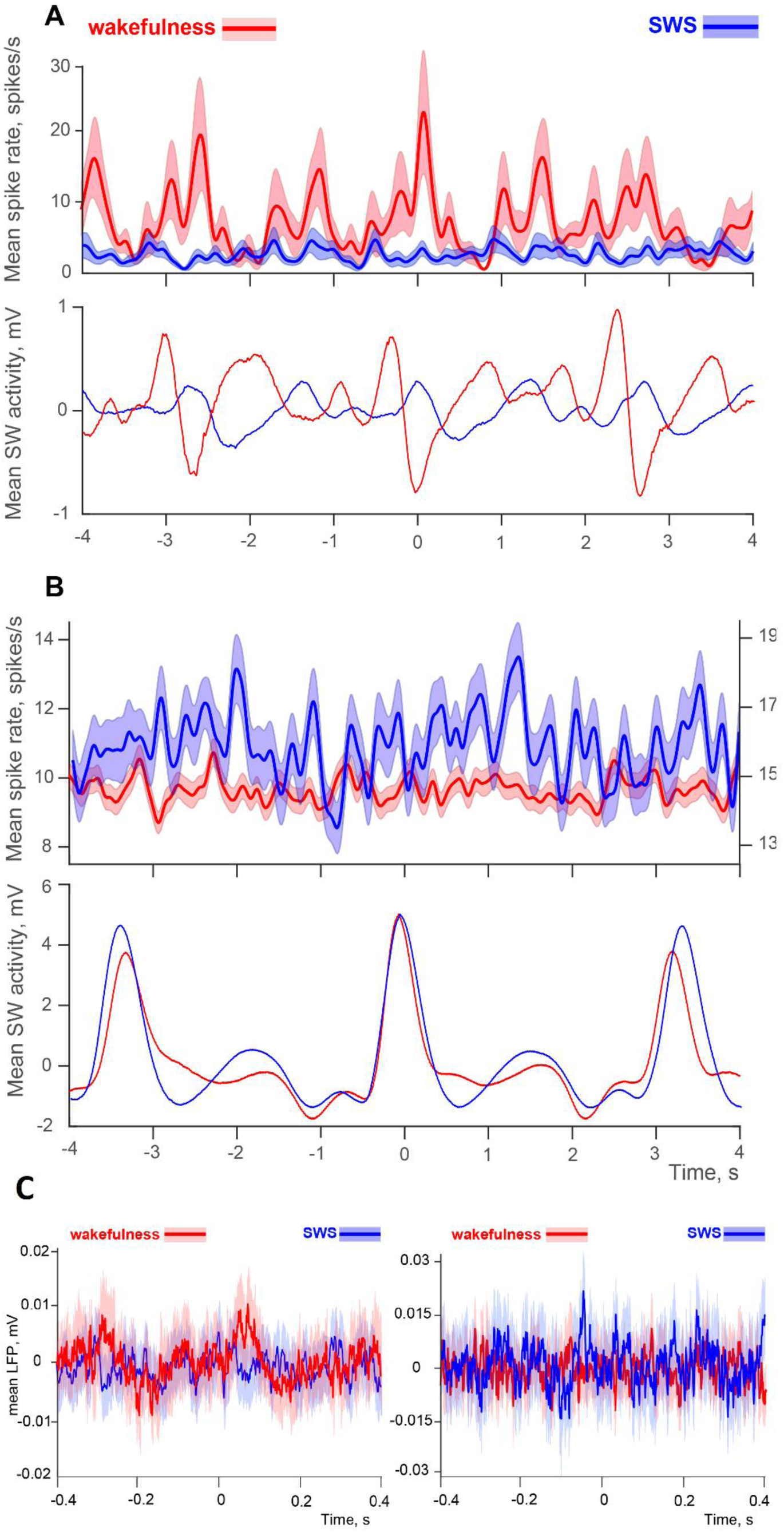
Synchronization of hippocampal activity with SW duodenal signals. A. Activities of a hippocampal cell synchronized to SW in wakefulness, top row. The corresponding averaged duodenal myoelectric activity signals are in the bottom row. B. Activity of a hippocampal cell synchronized to SW in sleep, top row. The corresponding averaged duodenal myoelectric activity signals are in the bottom row. C. Event-related hippocampal LFPs responses to duodenal cycle in either wakefulness (left side) or sleep (right side).

The duodenum has the highest density of nutrient-detecting vagal afferents in the gastrointestinal system (Jagger, et al., 1997; Brookes et al., 2013), and hippocampal cells can potentially be modulated by strong signals related to the chemical composition of food. We hypothesized that prolonged exposure of duodenal receptors to chyme/food bolus would lead to more robust chemoreceptive signals ascending via the vagus nerve, thus leading to more robust synchronization of hippocampal cells to SW during sleep. LFP was found to be synchronized with SW duodenal activity in 20% of cases during SWS, 22.5% during wakefulness and 0% in both states of vigilance (Fig. 7.C). These proportions were significant according to binomial enrichment test, and marginal after FDR (q = 0.056 for both states of vigilance).

#### 2. Stomach

We observed nearly equal proportions of significant STAs of stomach activity in SWS and wakefulness, namely 29% (31/107) of cells in wakefulness, 26.2% (28/107) of cells in SWS and 8.4% of cells in both (9/107), both surviving enrichment test and FDR at q<0.001. 28% of them had high modulation significance (p<0.01). Additionally, in 8.4% of cells, significant STAs were observed simultaneously for stomach activity and the envelope of duodenal SB activity.

Figure 8 illustrates the complexity of hippocampal activity associated with gastrointestinal motility. The top rows of panel A show examples of spiking activities for two hippocampal cells, N1 and N2, recorded simultaneously from left and right hippocampi. The purple line shows the corresponding duodenal activity, and the brown line shows stomach activity, all during wakefulness. One can see that cell N1 drastically decreased its spiking when stomach activity increased, and more pronounced SB activity occurred. In contrast, cell N2 had increased spiking in corresponding to stomach activation, and that spiking was of a bursting type, to an extent synchronized to duodenal SB. Statistical analysis of the N2 cell activity revealed that it indeed had both significant STA for stomach activity and significant synchronization to duodenal SB, both occurring in wakefulness (panels B and C). These kinds of relationships between hippocampal and visceral activities can potentially underlie effects such as eating epilepsy (Nagaraja and Chand, 1984; Girges et al., 2020), where strong gastrointestinal stimulation triggers epileptiform activity.

**Figure 8.**
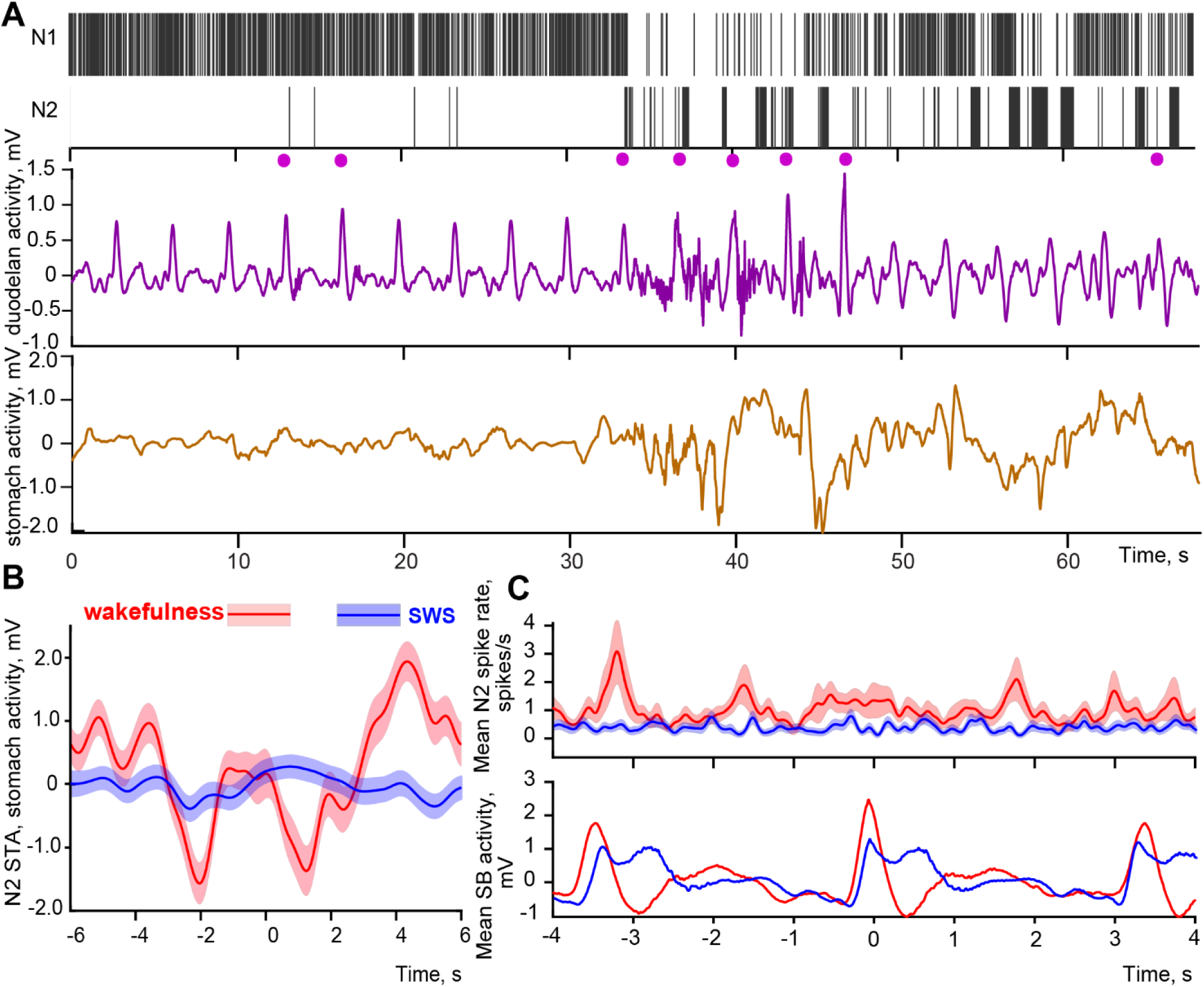
Hippocampal activity associated with gastrointestinal motility. A. Top two rows show examples of spiking activities for two hippocampal cells, N1 and N2, recorded simultaneously from left and right hippocampi during wakefulness. Purple trace in the next row below represents the corresponding co-registered duodenal activity, with purple dots marking SB waves, and the brown trace in last row represents the corresponding stomach activity. B. STA of stomach myoelectric signal to spiking of cell N2. C. Event-triggered responses of cell N2 to the peaks of SB waves of duodenal activity.

### SYNCHRONIZATION OF HIPPOCAMPAL CELLS TO MORE THAN ONE TYPE OF VISCERAL EVENTS

Hippocampal cells were only rarely synchronized with both cardiorespiratory and intestinal activity. Only 3.1% of cells that had synchronization to breathing were also synchronized to either SB or SW within the same state of vigilance, and just 0.9% of co-synchronization was observed between heartbeats and either SB or SW. Cardiac and respiratory activities are known to be highly correlated and the same can be said for gastric and duodenal activities. However, the relationships between the events belonging to these two different groups are not expected to be as strong (e.g., gastric and cardiac events are less interrelated than cardiac and respiratory ones). One can see that for these physiological events with weak functional relationships; hippocampal cells did not demonstrate co-synchronizations as well.

In contrast, the levels of conjoined modulation were high within breathing (when both event-triggered and spike-triggered analyses showed significant modulation for the same cell in the same state of vigilance), with co-modulation levels equal to 21.8%. Conjoined synchronization to breathing and heartbeats was at 5.9%. Additionally, 8.4% of cells had significant STAs for both stomach activity and the envelope of duodenal SB activity. Thus, co-activation was higher between functionally related signals compared to unrelated ones.

To summarise the results, Table 2 demonstrates percentages of hippocampal cells whose activities were significantly associated with different types of visceral signals, for pooled data (A) as well as individually for each animal (B).

**Table 2.** Relationships between recorded visceral signals and hippocampal activity. (A) Results are presented for both animals together as percentages of significant relationships between hippocampal and visceral signals in either wakefulness or in SWS (bootstrapping-based analysis, p<0.05). * indicates significant predisposition of visceral activity to be linked to hippocampal activity in one of the states of vigilance more frequently that in the other (χ2 with Yates correction, p<0.05).Spike-triggered results for the heart rate are excluded to avoid possible confound (see Results for further details, section STA OF THE HEART RATE). (B) Relationships between recorded visceral signals and hippocampal cells activity for each of two animals. * indicates a significant predisposition of that visceral activity to be linked to hippocampal cells activity in a particular state of vigilance (χ^2^ with Yates correction, p<0.05). For the details on the numbers of cells used see Materials and Methods, subsection titled NUMBER OF CELLS RECORDED AND AVAILABLE FOR THE ANALYSIS.

|  | Event-triggered analysis |  |  |  | Spike-triggered analysis |  |  |
| --- | --- | --- | --- | --- | --- | --- | --- |
| State of vigilance | Breathing, % and N cells | Heart beats, % and N cells | Duodenal SB, % and N cells | Duodenal SW, % and N cells | Breathing amplitude, % and N cells | Duodenal SB envelope, % and N cells | Stomach, % and N cells |
| Awake only | 20.1%<br>37/184 | 7.1%<br>15/213 | 10.8%<br>23/213 | 6.6%<br>14/213 | 23%<br>42/184 | 11.3%*<br>24/213 | 29%<br>31/107 |
| SWS only | 22.8%<br>42/184 | 8%<br>17/213 | 10.3%<br>22/213 | 15.5%*<br>33/213 | 42.9%**<br>79/184 | 6.15%<br>13/213 | 26.2%<br>28/107 |
| both | 3.8%<br>7/184 | 0%<br>0/213 | 0.95%<br>2/213 | 0.95%<br>2/213 | 12%<br>22/184 | 1.9%<br>4/213 | 8.4%<br>9/107 |

B.
|  |  | Event-triggered analysis |  |  |  | Spike-triggered analysis |  |  |
| --- | --- | --- | --- | --- | --- | --- | --- | --- |
| State of vigilance | Cat | Breathing, N cells | Heart beats, N cells | Duodenal SB, N cells | Duodenal SW, N cells | Breathing amplitude, N cells | Duodenal SB envelope, N cells | Stomach N cells |
| Awake only | **1** | 24/121 | 12/121 | 11/135 | 7/135 | 24/121 | 10/135 | 17/72 |
|  | **2** | 15/63 | 3/63 | 11/78 | 10/78 | 18/63 | 14/78 | 14/37 |
| SWS only | **1** | 27/121 | 6/121 | 10/135 | 14/135* | 48/121* | 6/135 | 22/72 |
|  | **2** | 13/63 | 12/63 | 13/78 | 18/78* | 31/63* | 7/78 | 6/37 |
| both | **1** | 6/121 | 0/121 | 1/135 | 1/135 | 13/121 | 0/135 | 7/72 |
|  | **2** | 1/63 | 0/63 | 1/78 | 1/78 | 9/63 | 4/78 | 2/37 |

Results of the LFP analysis demonstrate nearly equal proportions of sites responding in either wakefulness or in sleep in synchrony to duodenal SW and SB waves, with absence of simultaneous responses in both states. This indicates a high level of segregation among the networks involved in the analysis of intestinal information. Since LFPs are known to primarily reflect inputs to the network while spikes are output-related (Logothetis, 2002; Raichle and Mintun, 2006), the observed state-specificity in cell responses, combined with the separation of inputs, suggests that specific types of processing related to digestion occur in the hippocampus in both states, albeit in very different ways that may support different functions. Similar differential processing of interoceptive information was observed earlier in the insular cortex (Levichkina et al., 2021).

The signal that had a relatively high proportion of LFPs responsive in both states was breathing. However, Similarity index analysis (Fedorov et al., 2023) suggested preferential synchronization of the hippocampus to breathing during episodes of high power of delta waves in the EEG, which occurred not only during sleep but also during microsleep episodes in wakefulness. The LFP responses associated with cardiac cycle also frequently occurred in both states, but they had a predilection to synchronize at the presence of either spindles or delta waves even during wakefulness. This highlights the need for a more detailed analysis of LFP responses, especially when these responses appear to occur indiscriminately between sleep and wakefulness.

## Discussion

A growing corpus of literature indicates that bodily rhythms structure hippocampal activity during ongoing behaviour. Respiration-coupled hippocampal oscillations are present in awake rodents, they modulate neuronal activity and sharp-wave ripples (Nguyen Chi et al., 2016; Lockmann et al., 2016; Liu et al., 2017). In humans, neuronal firing in parahippocampal cortex covaries with cardiac-cycle duration, and breathing coordinates hippocampal sleep oscillations, further supporting the idea that visceral rhythms provide a temporally relevant organizing signal for medial temporal circuits across vigilance states (Kim et al., 2019; Sheriff et al., 2024). At the same time, the relationship is not strictly unidirectional, because hippocampal stimulation can alter heart rate, blood pressure, respiration, and gastric motility, indicating that hippocampus is embedded in reciprocal autonomic loops rather than passively entrained by them (Ruit and Neafsey, 1988; Guan et al., 2003). Against this background, the main advance of our dataset is the demonstration across multiple simultaneously monitored visceral modalities, that substantial synchronization between visceral and hippocampal activities is present in wakefulness and that sleep reshapes its distribution across signals and neuronal subpopulations. The rarity with which neurons coupled to one visceral modality in one vigilance state remain coupled to that same modality in the other state argues for dynamic state-dependent reconfiguration of hippocampal visceral coupling, rather than a fixed class of “visceral” cells. Our data emphasises persistent body-hippocampus synchronization in both wakefulness and sleep, with state-dependent enhancement for selected signals emerging on top of this broader and functionally important organization.

A substantial proportion of hippocampal cells in our study showed spike activity synchronized to either rhythmic visceral events or to pronounced arrhythmic ones. These cells could be modulated by both types of visceral activity (see Figs 2 and 4 for cardiorespiratory activity, and Fig 8 for gastrointestinal activity, and Figure 9 for an overview of the changes per state of vigilance). All event-triggered and spike-triggered measures for these cells showed statistical significance rates above the 5% error threshold, ranging from 6.6% to 42.9% of cells. Most comparisons had over 5% of cells showing statistical significance at p<0.01. To strengthen the analytic rigor of the results we have applied a double approach to tackling multiple comparisons, introducing binomial enrichment test followed by FDR. Significant relationships between visceral and hippocampal signals that can be reported after this analysis for both states of vigilance are all of the breathing-related and stomach-related ones. Synchronization between hippocampal cells spiking and duodenal SW was significant only in SWS. Duodenal SB and envelope of high-frequency duodenal signal only passed both the enrichment test and FDR in wakefulness. While heartbeats and hippocampal spiking synchronization did not pass our thresholds in either wakefulness or SWS, LFP synchronization was significant for both states. We therefore conclude that hippocampal activity is potentially linked to interoceptive signals.

**Figure 9.**
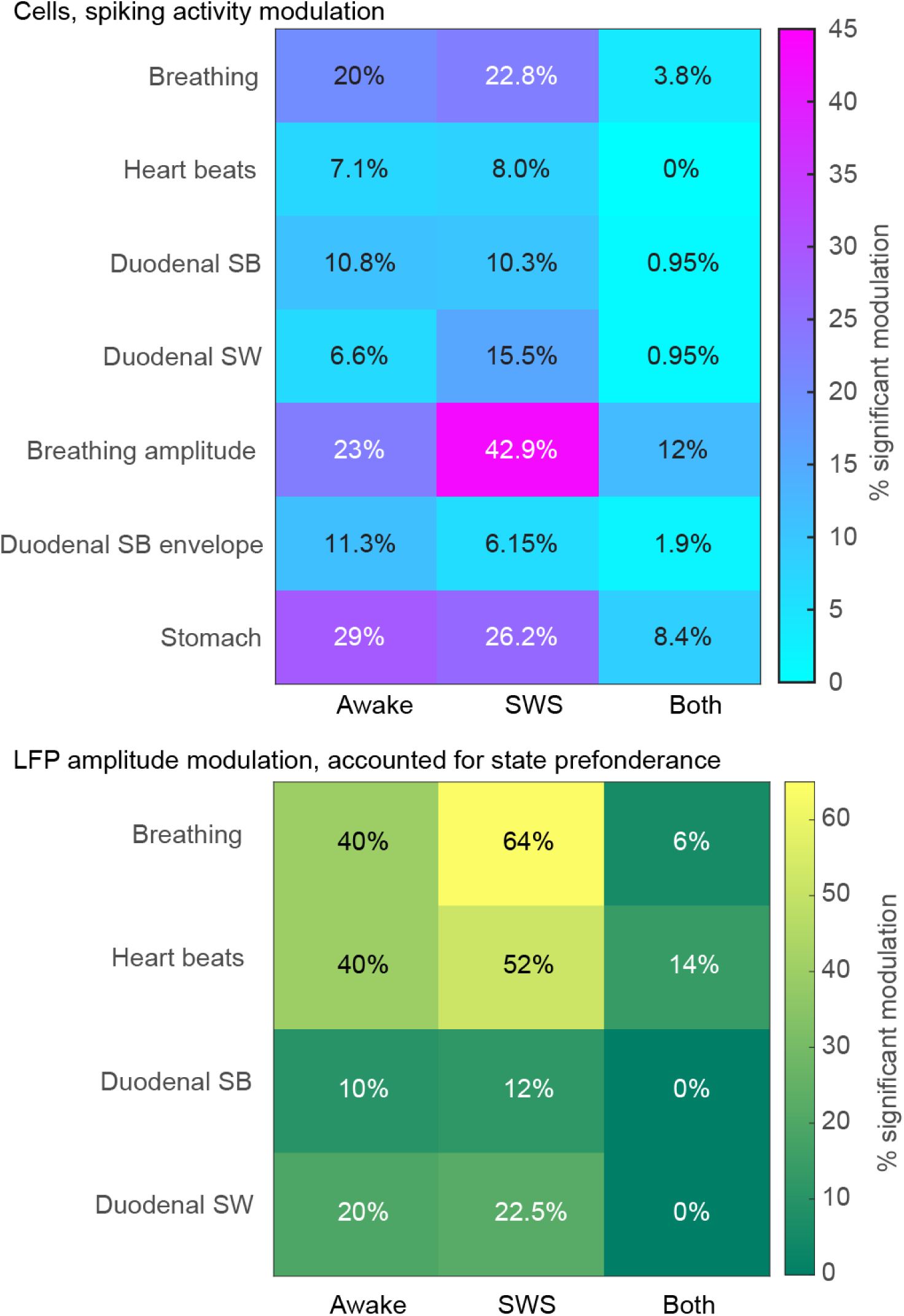
Overview of modulation of hippocampal activity during wakefulness, SWS and in both states. Heatmaps of percentages per state of vigilance are given for spiking activity (top panel) and LFPs (bottom panel).

The relationships between visceral and hippocampal activities could be significant in either wakefulness or sleep, but the majority of hippocampal cells synchronized only in one state. This was evident with all types of the visceral signals we studied. However, two of the calculated types of STAs showed dual (i.e., in SWS and wakefulness) preferences, namely in 8.4% of recordings from the stomach and in 12% of recordings of respiration. This may indicate that tracking rhythmic visceral events is more vigilance state-specific than tracking visceral events of a particular strength. However, the proportion of cells linked to these events was also the highest in our sample (23% in wakefulness and 42.9% in sleep for STA analysis), reflecting a potentially greater relationship hippocampal activity to the high amplitude visceral signals.

We observed co-synchronization of neuronal activities to functionally related visceral signals, such as cardiac and respiratory events (above the diaphragm) or duodenal and gastric (below the diaphragm), but not to the unrelated: i.e., activities of hippocampal cells could be linked to respiration and heart beats or to stomach and duodenal activities. These co-synchronizations we have observed might lead to the entrained frequencies of hippocampal cell spike trains having components distinct from events related to any one visceral system.

In line with previous research (Duffin and Hockman, 1972; Frysinger et al., 1989; Radna and MacLean, 1981; Poe et al., 1996; Herrero et al, 2018; Tort et al., 2018; Folschweiller and Sauer, 2021; Nokia and Penttonen, 2022), the strongest link was observed between respiration and hippocampal activities, particularly during sleep, with 42.9% of cells showing significant breathing STAs. Hippocampal LFPs also showed the highest rates of significant responses to breathing, especially during high delta power episodes in SWS and wakefulness. Thus, LFPs were also becoming synchronized to respiration in sleep or drowsiness.

Hippocampus is known to be involved in respiratory control, and its stimulation can alter breathing patterns, including evoking apnoea (Kaada and Jasper, 1952; Duffin and Hockman, 1972; Lacuey et al., 2017; Lacuey et al., 2019; Limanskaya et al., 2024). Our results provide the first direct neural evidence for reports of hippocampal involvement in high-amplitude respiratory events (Poe et al., 1996; Ajayi and Mills, 2017), but particularly during sleep. This may lead to oversynchronization of vulnerable hippocampal circuits during SWS, potentially explaining the link between sleep-disordered breathing and epilepsy (Harnod et al., 2017), as well as the induction of paroxysmal gamma waves in the temporal lobe by specific breathing practices (Vialatte et al., 2009). However, as our study does not allow to resolve the direction of the effects and demonstrate whether the driver of such synchronization comes from the visceral system or the hippocampus, further investigation of the directionality is necessary to establish any causal relationships here. In addition, our experiments were conducted in healthy animals, and an appropriate epilepsy model is required to study whether the interoceptive signalling is the one triggering oversynchronization. It is also possible that two systems are both prone to synchronization at similar frequencies and can mutually enhance these effects. At the same time, one may argue that the ascending driving force is more likely in the case of breathing. For example, Lockmann et al. (2016) showed that a hippocampal respiration-coupled rhythm was abolished by tracheotomy and restored by rhythmic nasal air puffs, which strongly supports an ascending drive rather than the hippocampus imposing the rhythm on the body. Nguyen et al. (2016) further reported that the hippocampal respiration-driven rhythm is distinct from theta and that directionality analysis is consistent with a drive from the olfactory bulb to hippocampus. In humans in was demonstrated that breathing organizes hippocampal sleep oscillations, again favouring an ascending respiratory influence on hippocampal synchronization (Sheriff et al., 2024).

Our results show moderate synchronization of hippocampal neurons to heartbeats with none significantly related to heartbeats in both states. Responses to cardiac events were reported before (Frysinger and Harper, 1989; McEchron et al., 2000), hippocampal stimulation also changes heart rate and blood pressure (Kaada and Jasper, 1952; Anand and Dua, 1956), and hippocampus is involved in controlling heart rate in humans (Norton et al., 2013).

High occurrence of significant heart rate STAs likely resulted from the vigilance state dependency of both hippocampal cell spiking and heart rate. Changes in heart rate across vigilance states can produce apparent state-dependent coupling even without a direct hippocampal influence, and emotional arousal can similarly co-modulate both variables. At the same time, human intracranial recordings show that resting firing fluctuations in medial temporal and cingulate regions can predict cardiac-cycle duration, with analyses favouring a neural-to-cardiac direction in many units (Kim et al., 2019). Therefore, our data are compatible with both shared-state co-modulation and a potential top-down hippocampal contribution to cardiac regulation, but do not by themselves disambiguate these mechanisms. However, this effect might explain the value of heart rate signals in seizure prediction (Van Elmpt et al., 2006; Sevcencu and Struijk, 2010; Fujiwara et al., 2016; Stirling et al., 2021). LFP responses showed higher occurrences and overlaps between vigilance states compared to cell responses; however, they demonstrated state preference as well, favouring episodes with either higher spindle activity or higher delta activity.

We observed 5.9% of cells synchronizing with both breathing and heartbeats in the same state of vigilance. Such co-modulation can further increase the chances of modulation of hippocampal activity by related cardiorespiratory events. Since LFP synchronization to breathing occurred nearly exclusively during periods of strong delta activity, while synchronization to heartbeats happened during high delta or high spindle activities, it seems that both visceral inputs were also more effective in synchronizing hippocampal activity during sleep or microsleep episodes.

Recent studies had uncovered a widespread brain network associated with stomach activity in the resting state (Rebollo et al., 2018) and substantial increase of brain responses to vagal (Rembado et al., 2021) and intestinal (Levichkina et al., 2021) stimulations in SWS. Our current study has provided the description of hippocampal synchronization to gastrointestinal activity at the neuronal level. Hippocampal activity was linked to duodenal and gastric myoelectric signals.

Higher frequency duodenal signals (SB envelope) which are indicative of duodenal motility were linked to hippocampal activity during wakefulness, while regular duodenal SWs synchronized with the hippocampus more effectively during sleep. LFP analysis showed distinct duodenal inputs to the hippocampus across different states of vigilance. Hippocampal sites synchronized with duodenal activity exclusively in either wakefulness or sleep, not both. This suggests a strict separation between the inputs into networks processing GI signals during sleep and wakefulness, similar to observations made for the insular cortex (Levichkina et al., 2021).

Certain GI-related signals are related to wakefulness, while the others are likely to occur more at rest. For example, peak gastrointestinal mechanosensitivity (Page, 2021) and related mRNA and protein production (Saito et al., 2008; Voigt, Forsyth, Keshavarzian, 2019) occur during wakefulness and the SB component of duodenal signals are more representative of these in comparison to SW (Papasova and Milenov, 1965; Papasova et al., 1966; Costa and Furness, 1982; Sarna, 1989; Martinez-de-Juan et al., 2000). In contrast, satiety causes postprandial sleepiness in many species, including humans (Antin et al., 1975; Borbély, 1977; Danguir et al., 1979; Orr et al., 1997; Wells et al., 1998; Shemyakin and Kapás, 2001; Bazar et al., 2004; Murphy et al., 2016) leading to rest or sleep. Postprandial somnolence is largely mediated by vagal signals related to gastric distension, food composition, and humoral influences (Hansen et al., 1998; Shemyakin & Kapás, 2001; Kim and Lee, 2009).

In line with the above as well as the food transition times (Camilleri, 2019; Husnik et al., 2017; Telles et al., 2022), a substantial part of digestion and the highest levels of nutrient absorption occur during resting periods or sleep, while the higher duodenal motility happens in awake state (Duboc et al., 2020). Satiety-related gastrointestinal signals also induce low-frequency EEG synchronization (Hansen et al., 1998). The increase in phase-amplitude coupling between MEG-measured brain activity and electrogastrographic signal was also observed during rest (Balasubramani et al, 2022). Our results for duodenal signal synchronization are consistent with the known differences of GI activity in different states of vigilance. However, analysed SW events represent ongoing duodenal electrical activity; we did not manipulate feeding state, nutrient composition, or luminal delivery, so nutrient dependence of the observed SW and hippocampal coupling cannot be directly concluded from our data. We therefore treat the SW link to nutrient absorption as a hypothesis which can be biologically plausible because nutrient exposure in the duodenum is known to generate nutrient-specific vagal afferent signalling and coordinated motility responses (Schwartz and Moran, 1998), and postprandial intestinal SW/SB organization can be enhanced through cholinergic mechanisms, as shown by atropine sensitivity (Qin et al., 1999). Thus, nutrient-vagal modulation is a reasonable candidate mechanism for future testing, but the present results should be interpreted as evidence for state-dependent SW and hippocampus synchronization rather than direct proof of a nutrient-driven pathway. Previous studies suggest that gastrointestinal influences are very likely to drive hippocampal activity. Suarez et al. (2018) showed that selective gastrointestinal vagal deafferentation impairs hippocampal-dependent memory and noted that hippocampal neurons could be activated by direct vagus nerve stimulation and by GI vagally mediated signals such as gastric distension and intestinal nutrient-related input. Consistent with this ascending interpretation of the driver of synchronization, Wang et al. found that activation of an implantable gastric stimulator increased metabolism in the hippocampus in humans (Wang et al., 2006). Taken together, these findings support the view that at least part of the coupling between GI and hippocampal activity can be driven from the periphery upward, while still not excluding descending or common driver mechanisms, considering, e.g., that motilin injection into the hippocampus can affect gastric motility (Guan et al., 2003).

8.4% of cells in our study exhibited significant STAs for both stomach activity and the duodenal SB envelope. Since increased stomach motility is likely associated with higher duodenal motility, this co-activation leading might be a common signal, which, provided that the direction of signalling is from the visceral system to the hippocampus, can be consistent with strong association between eating and seizures (Nagaraja and Chand, 1984; Tényi et al., 2021).

Neurophysiological studies involving large animal models commonly rely on data from a pair of animals with multiple neurons sampled from each of them. In this approach cells are recognised as the units of statistics and two animals are used to avoid a situation when a sample gets recorded from an animal that is by itself an outlier. However, it is important to acknowledge that our animal sample size is limited to only two cats and relatively small number of locations within dorsal hippocampus. Although our animals demonstrated comparable results, it is quite possible that the reported proportions of cell activities might be clarified by using more animals and different recording positions along the ventro-dorsal axis of the hippocampus. Our recordings were restricted to the dorsal hippocampus, while ventral/anterior sectors are often considered more tightly linked to viscerolimbic and epilepsy-relevant circuitry (Fanselow and Dong, 2011; Strange et al., 2014). This creates a risk of overgeneralizing dorsal findings to the full longitudinal axis of the hippocampus.

However, in cats, dorsal hippocampus itself is not functionally isolated from interoceptive processing: classic feline work shows graded dorsal–ventral differences in firing organization and projection architecture rather than a strict binary split (Elul, 1964a,b; Siegel and Tassoni, 1971). Our data extend this by showing dorsal hippocampal locking to multiple visceral signals (cardiac, respiratory, gastric, and duodenal) across vigilance states. At the same time, translational interpretation should remain cautious because human mesial temporal epilepsy is frequently centred in anteromesial temporal structures, with prominent involvement of hippocampal head and entorhinal circuitry (Engel, 2001; Bernasconi et al., 2003; Bartolomei et al., 2005).

Brain activity, including in the hippocampus, is more synchronized during SWS (Halgren et al., 1978; Ferrara et al., 2012) and epileptic discharges often depend on the same mechanisms as slow-wave activity (Beenhakker and Huguenard, 2009). While the sympathetic system supports requirements of wakefulness, parasympathetic system is more active during sleep (Trinder et al., 2001; Cabiddu et al., 2012; Fatt et al., 2020). Interoceptive inputs amplified by the parasympathetic system are therefore likely to be stronger during sleep. The vagus nerve is the major source of this input to various regions of the brain. Hippocampus receives polysynaptic vagal input via medial septum (Castle et al., 2005; Suarez et al., 2018), which is ideally positioned to regulate transmission of interoceptive signals to the hippocampus between states of vigilance (Levichkina et al., 2025). Many areas receiving vagal input are also connected to the hippocampus (Kishi et al., 2000; Krout et al., 2002; Saper et al., 2005; Castle et al., 2005; Herman et al., 2005; Dong and Swanson, 2006; Cenquizca and Swanson, 2007; Petrovich et al., 2001; Fanselow and Dong, 2011).

Both the input (LFP) and the output (neuronal spiking) signals demonstrated association with various visceral events. We suggest that the higher propensity for synchronization during SWS, coupled with sleep-linked visceral events such as breathing and SW may trigger paroxysmal activity (Levichkina et al., 2025). Conversely, desynchronized hippocampal activity during wakefulness may oppose such triggering.

To summarize, all the visceral signals we studied were reflected in hippocampal activity. Sleep not only did not prevent this synchronization, but for respiration and periodic duodenal activity, increased its probability.

## Acknowledgments

This project was supported by the Russian Foundation for Basic Research Grants and the National Health and Medical Research Council (Project No2027651). Ivan Pigarev is gratefully acknowledged by the other authors for his original contribution with conceptualization, collection of data and discussion of the results prior to his unfortunate death on 15 July 2021.

